# DNA Damage Response Proteins Are Involved in the Formation of Defective HIV-1 Proviruses

**DOI:** 10.64898/2026.03.31.715508

**Authors:** K. Michalek, S. Bhattacharjee, A. Movasati, V. Clerc, J. Andres, A. Hotz, D. Rodrigues, K.J. Metzner

## Abstract

Latent HIV-1 proviruses remain the major barrier to curing HIV infection. Although many of these proviruses are defective, with large internal deletions and hypermutations, the mechanisms underlying their formation are still poorly understood. In this study, we applied CRISPR/Cas9 knockout screens to identify DNA damage response (DDR) proteins that contribute to the formation of defective HIV-1 proviruses carrying large internal deletions. Using an HIV-1-based dual-fluorophore vector as a model, we distinguished cells harbouring intact proviruses from those carrying large internal deletions by flow cytometry and cell sorting. We then validated top candidates using CRISPR-mediated gene activation and small interfering RNA-mediated knockdown.

Across these approaches, the helicase-like transcription factor HLTF emerged as a consistent modulator of large internal deletions: increased HLTF expression raised the proportion of cells carrying defective proviruses, whereas reduced HLTF expression had the opposite effect. Additional repair factors, including RAD1, RAD18, TREX2, and ZRANB3, also influenced the balance between intact and defective proviruses, suggesting that multiple DNA repair pathways cooperate in this process. Our data indicate that several DNA damage response proteins, including HLTF, are involved in the generation of defective proviruses and may constitute a previously undescribed host defence mechanism against HIV-1.

## Introduction

Human immunodeficiency virus type 1 (HIV-1) has caused more than 44 million deaths as the etiological agent of acquired immunodeficiency syndrome (AIDS) [1], and an estimated 40.8 million people were living with HIV-1 (PWH) at the end of 2024 [2]. Antiretroviral therapy (ART) effectively suppresses HIV-1 replication and reduces virus transmission; however, the persistence of the HIV-1 reservoir prevents cure of infection [3, 4].

A hallmark of HIV-1 infection at the cellular level is the integration of the viral genome into host chromosomal DNA, a process catalysed by the viral integrase [5]. The integration reaction leaves two single-stranded DNA gaps flanking the integration site, which are recognized as DNA damage and repaired by host DNA damage response (DDR) pathways [5, 6]. The precise mechanisms of post-integration repair remain incompletely understood, but several DDR pathways, including non-homologous end joining (NHEJ), base excision repair, and the Fanconi anemia pathway, have been implicated in this process [7–9]. Once repaired and integrated, proviruses can persist in a subset of cells and form the latent HIV-1 reservoir.

However, the majority of the integrated HIV-1 proviruses are defective in PWH on ART, with up to 98% defective proviruses detected in the CD4+ T cells from treated PWH [10–13]. The most frequent defects are large internal deletions (LID), which account for approximately 30-95% of all defective proviruses [10–13]. These deletions vary in size and position, with many proviruses carrying 3′ deletions affecting *env*, *tat*, *rev*, and *nef*, 5′ deletions affecting *gag* and *pol,* or very large deletions (> 6 kb) that remove most of the viral genome [10–13].

LID are assumed to arise from errors during reverse transcription, supported by the presence of short direct repeats present at some deletion junctions [10, 12, 14, 15]. Yet, in the study by Imamichi et al., such repeats were detected in only ∼40% of deletion junctions, leaving the mechanisms underlying the majority of LID unexplained [16]. Moreover, previous work from our group showed that LID occur in HIV-1-based vectors and in HIV-1 sequences integrated via CRISPR/Cas9, i.e. in the absence of viral reverse transcriptase and integrase [17, 18]. Together, these findings implicate the presence of additional host factors involved in the formation of LID. We hypothesize that these are most likely DDR proteins that are triggered during HIV-1 integration.

Here, we applied a targeted CRISPR/Cas9 screen to investigate whether DDR proteins contribute to the formation of defective HIV-1 proviruses carrying LID. As a screening readout, we used the HIV-1-based dual-fluorophore vector LTatC[M]L, which distinguishes double-positive Cerulean+/mCherry+ cells (integrated intact vector) from single mCherry+ cells (integrated vector with LID). In our previous studies, we have shown that the Cerulean expression could be reactivated only in a minor fraction of mCherry+ cells (<10%) following treatment with latency-reversing agents (TNF-α and SAHA, or TNF-α and Romidepsin) [17, 18]. Moreover, almost all non-inducible single mCherry+ cells carried an integrated defective vector (25/26 cell clones), containing LID in the HIV-1 Tat and/or Cerulean cassette, as opposed to double-positive Cerulean+/mCherry+ cells [17, 18]. By comparing guide RNA (gRNA) distributions between these populations, we generated a ranked list of host factors potentially involved in LID formation. Using a CRISPR activation (CRISPRa) system, we further showed that upregulation of several DDR proteins, including HLTF, increased the frequency of LID. These findings were corroborated by RNA interference-mediated gene silencing, where knockdown of *HLTF* alone, and in combination with *RAD1*, significantly reduced the number of LID. Together, our data support the involvement of multiple DDR proteins in the formation of defective proviruses, with implications for lentiviral gene delivery and HIV-1 cure strategies.

## Results

### Study design of the CRISPR/Cas9 DNA damage response (DDR) screening to identify genes potentially involved in the formation of large internal deletions (LID) in proviruses

We aimed to identify host factors that contribute to the formation of large internal deletions (LID) in HIV-1 proviruses. To this end, we performed a CRISPR/Cas9 knockout screen targeting 365 genes with known or suspected roles in the DNA damage response (DDR) as well as control core-essential and non-essential genes (Supplementary Table S1) [19]. The screening workflow comprised three main steps: transduction with a CRISPRko DDR library, transduction with the LTatC[M]L vector, and next-generation sequencing (NGS) readout (Figure 1a). First, SupT1 cells were transduced with a lentiviral CRISPRko DDR library. After puromycin selection, the resulting SupT1-CRISPRko DDR cells were transduced with the HIV-1-based dual-fluorophore vector LTatC[M]L. LTatC[M]L encodes mCherry, constitutively expressed from a human promoter, and Cerulean, expressed from the HIV-1 5’ LTR promoter [18]. Based on mCherry and Cerulean expression, cells were sorted into two populations: double-positive Cerulean+/mCherry+ cells harbouring integrated intact vector, and single mCherry+ cells, which have been shown to be non-inducible in our previous studies and carry defective vector with LID in the HIV-1 cassette [17, 18] (Supplementary Figure S1a). We expected gRNAs targeting DDR genes involved in LID formation to be under-represented in the single mCherry+ population, enriched for defective vectors, relative to the double-positive Cerulean+/mCherry+ population that contains intact vectors (Figure 1a). Genomic DNA from each of these cell populations was processed for NGS, and gRNA distribution was analysed using the MAGeCK (Model-based Analysis of Genome-wide CRISPR/Cas9 Knockout) pipeline [20] (Supplementary Figure S1b). In total, we obtained NGS data from three biological replicates of the DDR CRISPR/Cas9 screen (DDR1-3), each replicate comprising up to two independent NGS library preparations and two sequencing runs (Figure 1b).

**Figure 1.**
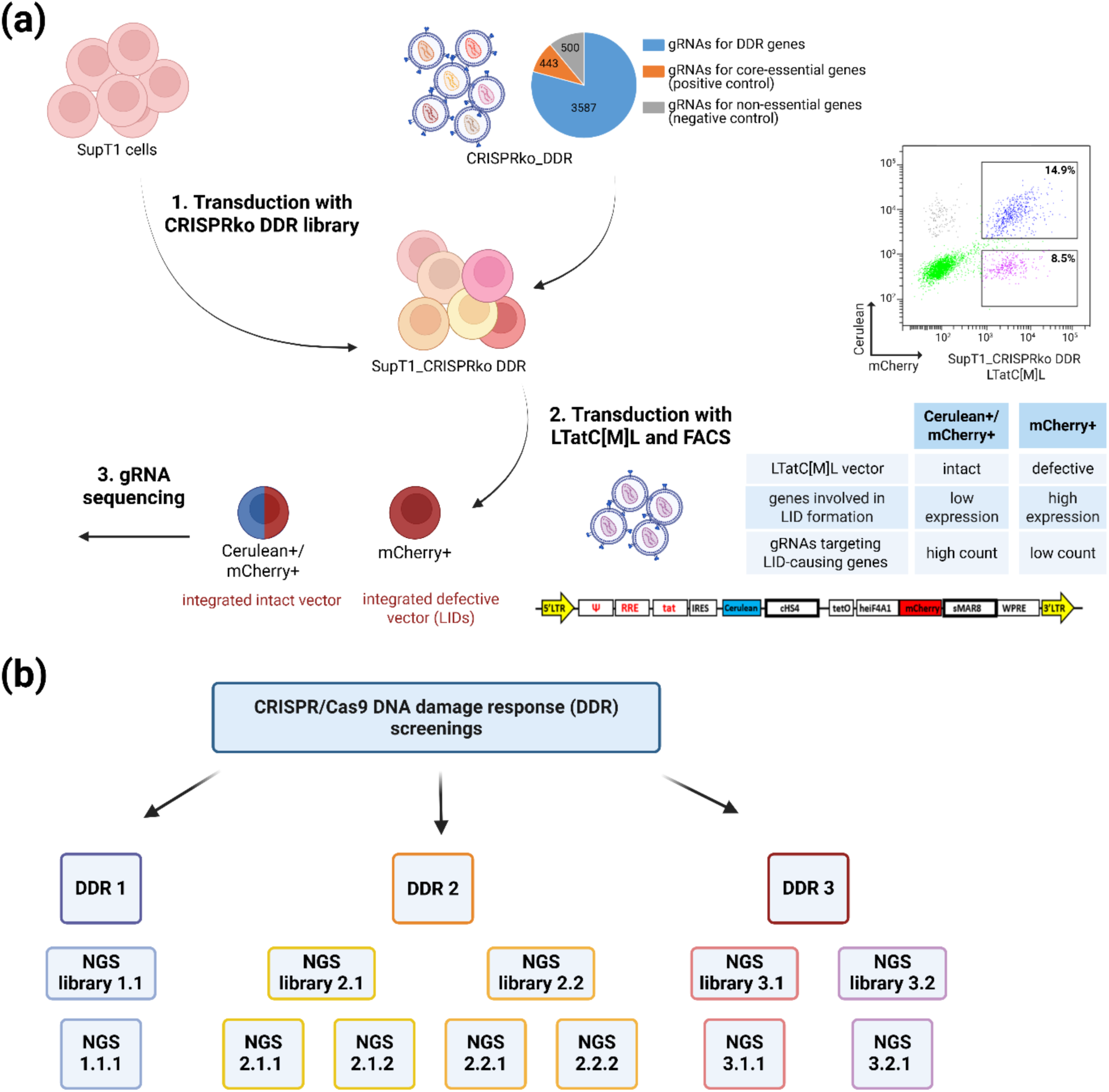
CRISPR/Cas9 DNA damage response (DDR) screening identifies genes potentially involved in the formation of large internal deletions (LID) in proviruses. (a) Schematic overview of the CRISPR/ Cas9 DDR screening workflow. Details are described in the text. (b) Overview of the CRISPR/Cas9 screening datasets. Abbreviations: *cHS4*, chicken hypersensitive site 4; *heIF4A1*, human eukaryotic initiation factor 4A1 gene promoter; *IRES*, internal ribosome entry site; *RRE*, HIV-1 *rev* responsive element; *sMAR8*, synthetic matrix attachment region 8 (from the human β-interferon gene); *tat*, HIV-1 transactivator; *tetO*, tetracycline operator sequences; *WPRE*; woodchuck hepatitis virus posttranscriptional regulatory element. Created with Biorender.

### Validation of the CRISPR/Cas9 DNA damage response (DDR) screening reveals a successful screening procedure

To minimize bias in the CRISPR/Cas9 screening, we first sequenced the CRISPRko DDR library after amplification. Using the approach described by Inderbitzin and Loosli et al., we confirmed that gRNA complexity was maintained in the library (Figure 2a) [21]. We observed ≤10% overlap in gRNAs between three independent sequencing samples and a similar number of observed and simulated unique gRNA sequences, indicating good gRNA representation in the library.

**Figure 2.**
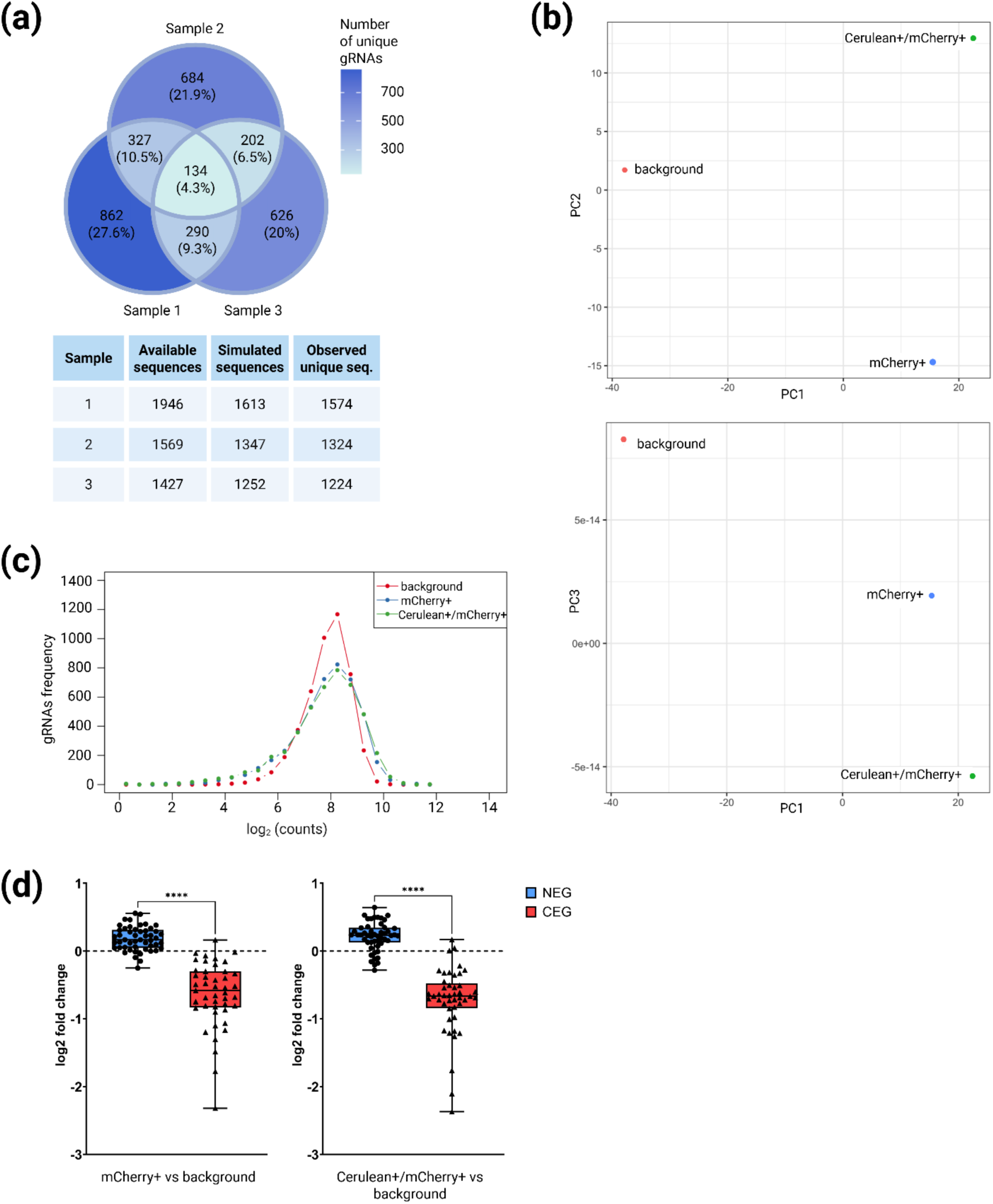
Validation of the CRISPR/Cas9 DNA damage response (DDR) screening. (a) Complexity estimation of the CRISPRko DDR library prior to the screening. From the top: overlap between gRNAs detected in three independent sequencing samples; Comparison between available and simulated number of unique gRNAs in the sequencing samples. (b) Principal component analysis (PCA) of the double-positive Cerulean+/mCherry+ cells, single mCherry+ cells, and background control samples. (c) gRNAs read count distribution in the double-positive Cerulean+/mCherry+ cells, single mCherry+ cells, and background control samples. (d) Internal controls of the CRISPR/Cas9 screening: log fold change in the count of gRNAs targeting core-essential (CEG) and non-essential genes (NEG) during CRISPR screening. Example plots shown for the second screening replicate (DDR 2). Data was statistically tested using a Mann-Whitney U-Test, p value <0.05 was considered significant, **** = p <0.0001. Created with Biorender.

After completing the CRISPR/Cas9 DDR screening, we assessed overall sequencing quality (Supplementary Table S2). Metrics included the number of mapped reads, the fraction of missing gRNAs, and the Gini index. gRNA read-count coverage ranged from at least 35x (DDR1) up to 1328x (DDR3). The proportion of missing gRNAs varied from 0.1-4% in the background control samples (prior to LTatC[M]L transduction) and from 0.1-20.6% in the sorted double-positive Cerulean+/mCherry+ and single mCherry+ samples. Importantly, the Gini index, which measures evenness of the gRNA read-count distribution, remained low: below 0.23 in background controls and below 0.42 in sorted samples, consistent with recommended thresholds [22].

Principal component analysis (PCA) showed clear separation of samples, with double-positive Cerulean+/mCherry+ and single mCherry+ populations clustering closer together compared to the background control, as expected (Figure 2b, Supplementary Figure S2a). The overall gRNA read-count distributions were largely similar between background, double-positive Cerulean+/mCherry+, and single mCherry+ samples (Figure 2c, Supplementary Figure S2b). As an internal control for screen performance, we compared counts of gRNAs targeting core-essential genes (CEG) and non-essential genes (NEG) (Figure 2d, Supplementary Figure S2c) [19]. The number of NEG-targeting gRNAs remained relatively stable across samples, whereas CEG-targeting gRNAs were significantly depleted in the single mCherry+ and double-positive Cerulean+/mCherry+ samples compared with the background control (p<0.0001), indicating effective negative selection and overall successful screening execution.

### Selection of the candidate genes involved in the formation of large internal deletions (LID) in proviruses

We next used (MAGeCK Model-based Analysis of Genome-wide CRISPR/Cas9 Knockout) to compare gRNA counts between double-positive Cerulean+/mCherry+ and single mCherry+ samples (Supplementary Table S3, Supplementary Figure S1b). Based on MAGeCK scores and associated fold changes, we identified 52 genes that were significantly enriched in the single mCherry+ population and therefore potentially involved in LID formation (Figure 3). Candidate selection began with the top 20 ranked genes (adjusted p ≤ 0.1) present in at least two technical replicates of the screen (Figure 1b), as summarized in Figure 3a. The list was then expanded by including the top 100 genes present across all three biological replicates (Figure 3b), resulting in a final set of 52 candidate genes (3 genes overlapping between Figure 3a and 3b). These genes were part of several DDR pathways, including NHEJ, homologous recombination, nucleotide excision repair, and the Fanconi anemia pathway. Notably, more than half of them have reported interactions with HIV-1 (Figure 3c).

**Figure 3.**
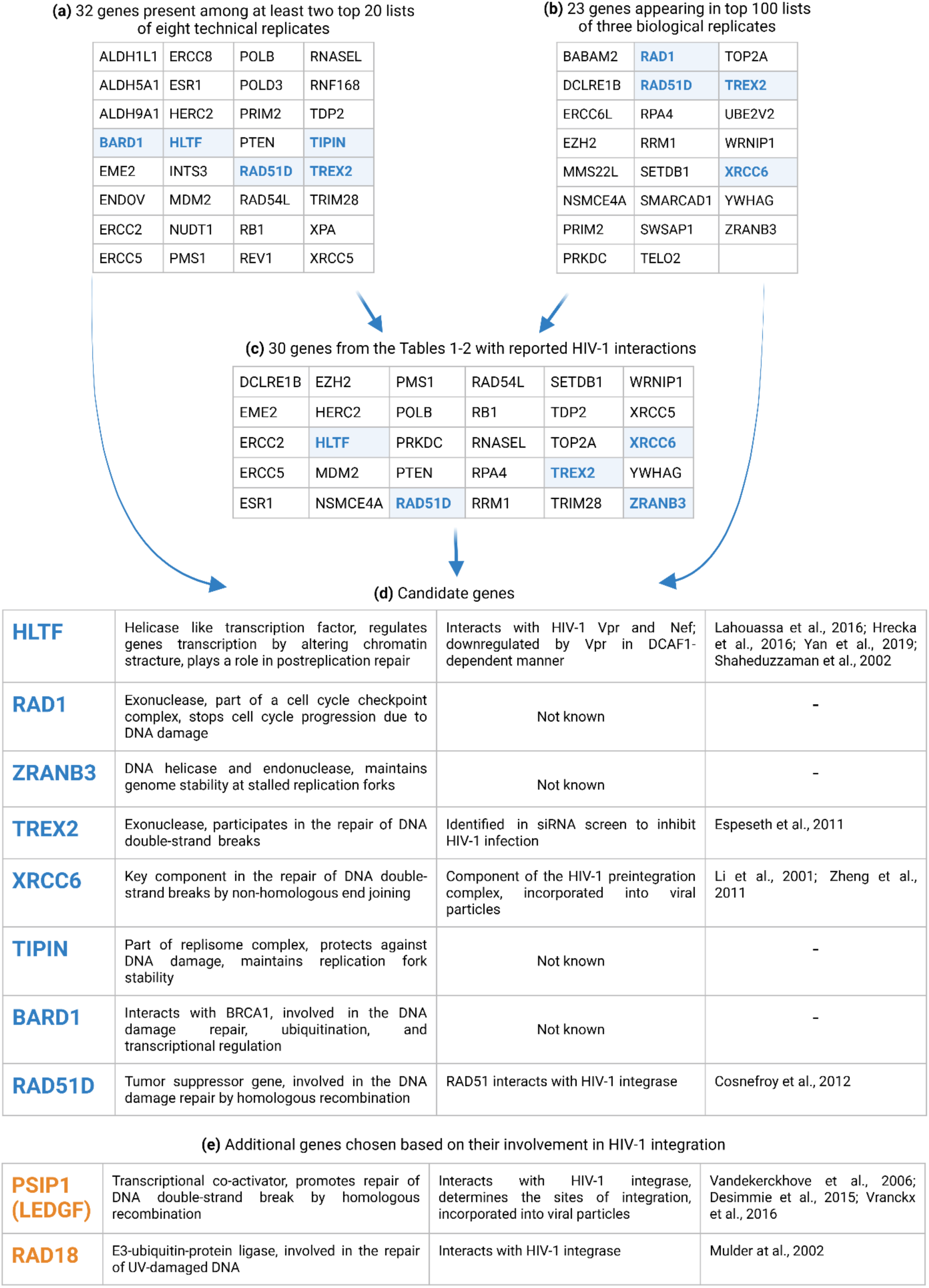
Selection of the candidate genes involved in the formation of large internal deletions (LID) in proviruses. The chosen candidate genes from the CRISPR/Cas9 DDR screening are highlighted in blue. Details are described in the text. Created with Biorender.

We selected eight genes for functional validation of their role in the formation of defective proviruses (Figure 3d). Two of them, *HLTF* (helicase-like transcription factor) and *XRCC6* (X-ray repair cross-complementing protein 6), have been reported to interact with HIV-1 proteins [23–27], whereas *TREX2* (three prime repair exonuclease 2) and *RAD51D* (RAD51 paralog D) have been indirectly linked to HIV-1 [8, 28]. The remaining four genes, *RAD1* (RAD1 checkpoint DNA exonuclease), *ZRANB3* (zinc finger RANBP2-type containing 3), *TIPIN* (TIMELESS-interacting protein), and *BARD1* (BRCA1-associated RING domain 1), have no reported HIV-1 interactions and were chosen solely based on their high MAGeCK ranking. In addition, we included two factors known to interact with HIV-1 integrase, *RAD18* (RAD18 E3 ubiquitin protein ligase) and *PSIP1* (PC4 and SRSF1 interacting protein 1; *LEDGF,* lens epithelium-derived growth factor), the latter of which was not part of the original CRISPRko DDR library (Figure 3e) [29, 30]. We measured basal mRNA expression of all ten candidate genes in SupT1 and K562 cells and observed low to moderate expression compared with *GAPDH* (Supplementary Figure S3a). Interestingly, all selected genes were reported to be downregulated at the mRNA level 12-18 h after HIV-1 infection in the study by Golumbeanu et al. Additionally, HLTF protein levels decreased as early as 6 h post-infection and remained reduced for up to 24 h (Supplementary Figure S3b) [31].

### DNA damage response (DDR) genes are involved in the formation of large internal deletions (LID) in proviruses

We next validated the role of the selected genes in LID formation using CRISPRa-mediated overexpression and small interfering RNA (siRNA)-mediated gene silencing approaches. K562-VPR cells stably expressing a dCas9-VPR cassette (nuclease-deficient Cas9 fused to VP64, p65, and Rta) were used for CRISPR-based gene activation with gene-specific gRNAs [32]. This complex acts in tandem to recruit transcriptional machinery and enhance transcription of target loci. Cells were transduced with lentiviral vectors encoding gene-specific gRNAs. For each gene, three different gRNAs were selected from commercially available CRISPR activation libraries (SAM and Calabrese) based on predicted off-target activity, self-complementarity, and binding efficiency [33, 34]. Titin (*TTN*) was included as a positive control because of its very low basal expression and robust detectability upon overexpression.

For each gene, the three gRNAs were cloned individually into the CRISPRa expression vector LentiGuide-Puro-P2A-EGFP, pooled, and packaged into lentiviral particles. K562-VPR cells were transduced at a multiplicity of infection (MOI) of 0.5, and GFP+ cells were isolated by FACS, indicating successful integration of gRNA constructs. GFP+ populations were subsequently expanded, and GFP expression remained stable at approximately 70% - 95% over time (Supplementary Figure S4a). These cell populations therefore expressed both the gRNAs and the dCas9-VPR activator, leading to upregulation of the corresponding target genes. Except for *RAD1* and *BARD1*, we confirmed moderate to high overexpression of all target genes at the mRNA level (Supplementary Figure S4c). Considering that GAPDH expression can vary depending on the cell activation state, expression was also normalized to *β-actin* for a subset of genes, yielding similar trends (Supplementary Figure S4c) [35]. *TTN* showed an approximately 75-fold increase in expression compared with the control, indicating efficient activation by the CRISPRa system. *TREX2* was not detectable by RT-qPCR, likely due to its extremely low basal expression.

CRISPRa-gRNA K562-VPR cells (GFP+) were then transduced in triplicate with LTatC[M]L at MOI 0.5. On day 7 post-transduction, flow cytometry was performed to quantify double-positive Cerulean+/mCherry+ and single mCherry+ cells using the gating strategy shown in Figure 4a. Observed LTatC[M]L transduction efficiencies were largely similar between the K562 VPR cells overexpressing target genes and non-targeting control gRNAs (Supplementary Figure S4b).

**Figure 4.**
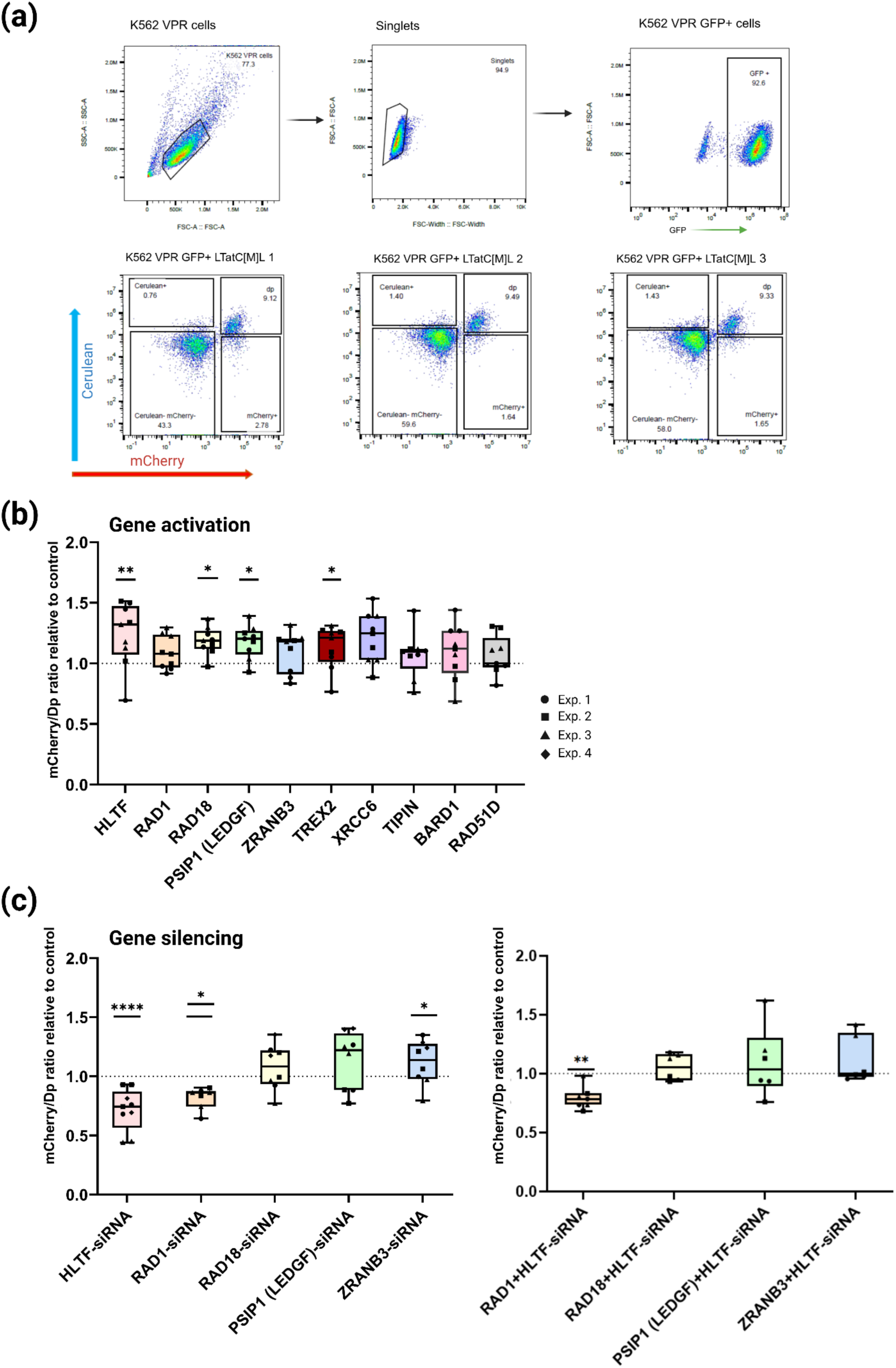
DNA damage response (DDR) genes are involved in the formation of large internal deletions (LID) in proviruses. (a) Flow cytometry gating strategy for K562 VPR cells transduced with CRISPRa-gRNA and LTatC[M]L lentiviral vectors. Example shown for CRISPRa-HLTF technical replicates. (b) Single mCherry+/double-positive Cerulean+/mCherry+ (mCherry/Dp) cell ratio for candidate genes in K562-VPR cells transduced with corresponding CRISPRa-gRNA and LTatC[M]L. Data shown relative to non-targeting control gRNA, marked by the dotted line. Each dot represents one technical replicate. (c) mCherry/Dp ratio for candidate genes in K562 cells transfected with corresponding siRNA and transduced with LTatC[M]L. From the left: transfection with siRNAs targeting single genes; transfection with a combination of siRNAs targeting two genes. Data shown relative to scrambled control siRNA, marked by the dotted line. Each dot represents one technical replicate. All data was statistically tested using a mixed-effect logistic regression model, p value <0.1 was considered significant. **** = p <0.001, *** p <0.01, ** p <0.05. Created with Biorender.

For each gene, raw cell counts in each population were used to calculate the change in the ratio of single mCherry+ cells to double-positive Cerulean+/mCherry+ cells, relative to cells transduced with non-targeting gRNAs. If a given gene contributes to LID formation, its overexpression is expected to increase this ratio, indicating a higher proportion of integrated defective vectors. To assess the effect of gene overexpression on this ratio, we applied a mixed-effects logistic regression model. Multiple-testing correction was performed using the Benjamini-Hochberg method; raw p-values ≤ 0.05 were considered statistically significant, and, in addition, adjusted p-values ≤0.1 were considered significant in this exploratory analysis. We used this relatively permissive adjusted-p threshold to reduce the risk of missing modest but potentially biologically relevant effects (Type II error), given the small sample size and variability (Supplementary Table S4).

Among the candidates, *HLTF* showed the most robust effect, with adjusted p = 0.03. Overexpression of *HLTF*, although modest in magnitude at the mRNA and protein level (Supplementary Figure S4c, S4d), consistently increased the proportion of single mCherry+ cells relative to double-positive Cerulean+/mCherry+ cells compared with the non-targeting control (Figure 4b). In addition to *HLTF*, *TREX2*, *RAD18*, and *PSIP1 (LEDGF)* also showed a significant increase in the single mCherry+ population, with raw p values < 0.05 and adjusted p values < 0.1 (Figure 4b). Protein overexpression relative to the control was verified by Western blot for a subset of genes, including those that showed a significant increase in the single mCherry+ population (Supplementary Figure S4d). Except for RAD1, proteins such as HLTF, RAD18, PSIP1 (LEDGF), TREX2, as well as ZRANB3 showed a modest increase in protein levels compared with the control.

Based on the gene overexpression results, we focused further analyses on five genes: *HLTF*, *RAD18*, *PSIP1 (LEDGF)*, *ZRANB3*, and *RAD1*, the last of which could not be efficiently overexpressed in the CRISPRa system. Despite promising CRISPRa results, *TREX2* was excluded from knockdown experiments because of its extremely low basal expression in K562 cells. Expression of the chosen genes was then silenced using an siRNA-based approach, with pools of three siRNAs targeting each gene. For all selected siRNA-targeted genes, we confirmed at least a 50% reduction at the mRNA level, with *HLTF* consistently showing the strongest knockdown (Supplementary Figure S5b-c). Protein expression was also reduced for all genes to approximately 0.6-0.8 relative to the control, except for HLTF, where the protein expression was so low it became undetectable (Supplementary Figure S5d).

Following the siRNA transfection, we chose the LTatC[M]L transduction time point individually for each gene based on siRNA kinetics, reported protein half-lives, and the estimated HIV-1 integration time of ∼15 h. This way, we could ensure the lowest gene expression during vector integration, when we expect the LID to be formed. mCherry and Cerulean expression was then measured 7 days post-transduction (MOI 0.5). Transduction efficiency was comparable between all siRNA-transfected samples, regardless of whether targeting siRNA or control siRNA was used (Supplementary Figure S5a).

To assess the effect of gene silencing on the ratio of single mCherry+ to double-positive Cerulean+/mCherry+ cells, we applied the same mixed-effects logistic regression model used for the CRISPRa experiments, again considering adjusted p values ≤ 0.1 as significant. If a chosen gene is involved in LID formation, upon its downregulation, we expected to see a decrease in the ratio of single mCherry+ cells to double-positive Cerulean+/mCherry+ cells, indicating a decrease in the number of integrated defective vectors.

We observed that siRNA-mediated silencing of *RAD1* and *HLTF* significantly decreased the ratio of single mCherry+ cells to double-positive Cerulean+/mCherry+ cells compared with control siRNA (*RAD1* adjusted p = 0.06; *HLTF* adjusted p < 0.0001; Figure 4c, left). This effect was maintained when *RAD1*- and *HLTF*-targeting siRNAs were co-transfected together (adjusted p = 0.03), supporting a role of both genes in the formation of LID in proviruses (Figure 4c, right). Downregulation of the other tested genes, *RAD18*, *PSIP1 (LEDGF)*, and *ZRANB3*, did not significantly affect the mCherry+ to double-positive cells ratio, except for *ZRANB3* (adjusted p = 0.08), where we observed a trend towards an increased proportion of mCherry+ cells (Figure 4c). Considering both CRISPRa-mediated gene overexpression and siRNA-mediated gene silencing experiments, *HLTF* emerges as the only gene that significantly affected the proportion of mCherry+ cells, containing LID.

### Large internal deletions (LID) are present in single mCherry+ cells

Our previous work showed that single mCherry+ cells predominantly harbour defective vectors with LID, whereas double-positive Cerulean+/mCherry+ cells contain largely intact vectors, serving as proxies for defective and intact HIV-1 proviruses, respectively [17, 18]. To confirm this and accurately map the deleted regions, we sorted single mCherry+ and double-positive Cerulean+/mCherry+ cells from LTatC[M]L-transduced GFP+ K562-VPR cells overexpressing *HLTF* (CRISPRa-HLTF), as well as from cells transduced with non-targeting control gRNAs (CRISPRa-NC), and LTatC[M]L-transduced SupT1 cells. The SupT1 cells were additionally treated with latency-reversing agents (TNF-α and SAHA) to re-activate single mCherry+ cells prior to sorting.

Genomic DNA was isolated from single mCherry+ cells and subjected to purification by pulsed-field gel electrophoresis (PFGE) to remove episomal forms and small DNA fragments (such as 2-LTR circles or unintegrated vector DNA), thereby enriching for high-molecular weight cellular DNA containing integrated proviral sequences. PFGE was performed using a BluePippin system, which is used for size selection of high-molecular weight DNA [36]. To assess the efficiency of plasmid DNA removal, we used SupT1 genomic DNA spiked with linear or circular forms of the HIV-1 all-in-one standard plasmid [37]. After PFGE, we observed a significant reduction in *pol* copy number (p = 0.002), indicating efficient depletion of plasmid DNA (Figure 5a left). Consistently, agarose gel electrophoresis of DNA before and after PFGE showed enrichment of high-molecular weight nuclear DNA and loss of lower-molecular weight episomal fragments and DNA smears (Figure 5a right).

**Figure 5.**
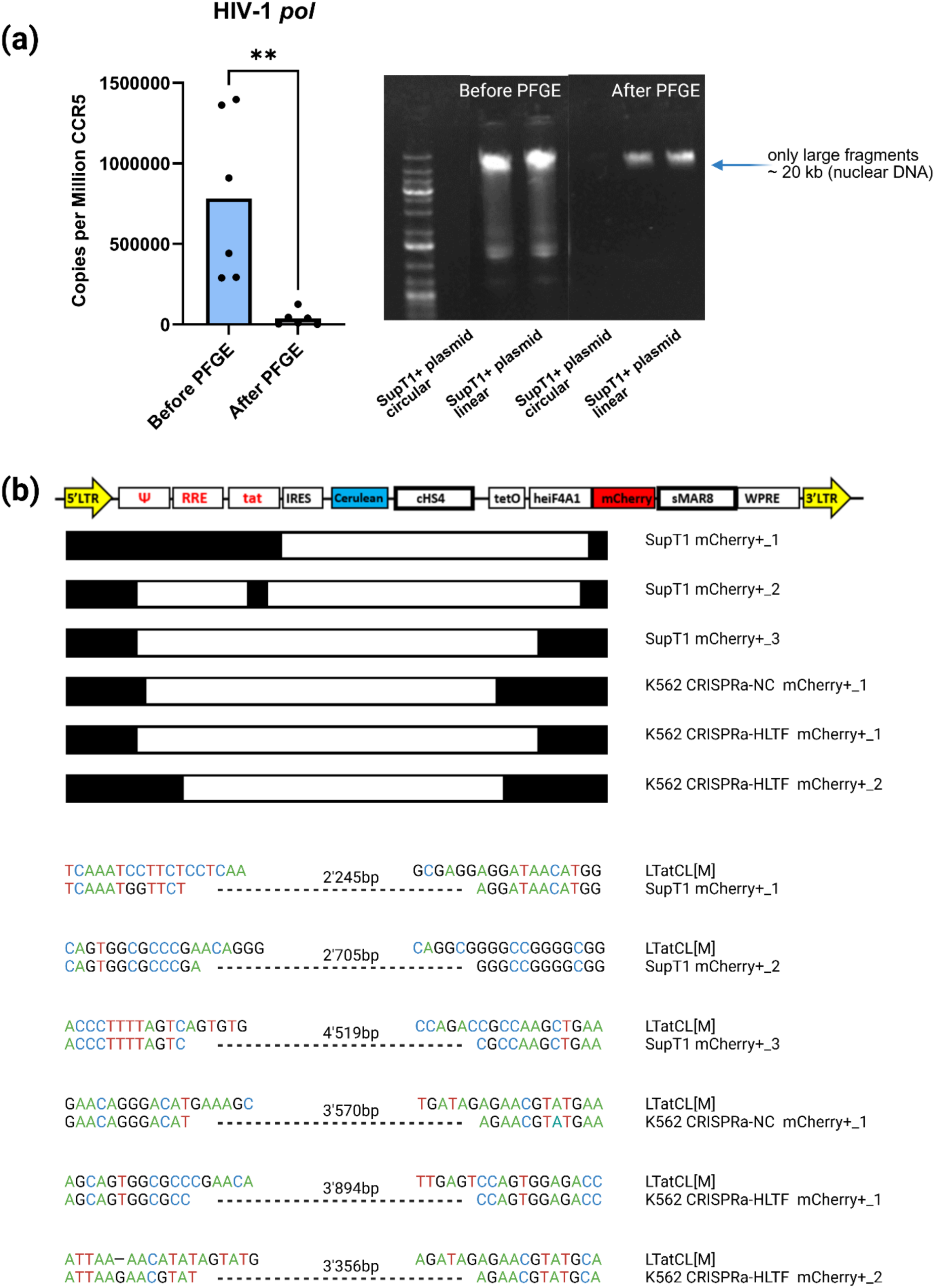
Large internal deletions (LID) are present in the mCherry+ cells. (a) Validation of the Pulse-Field Gel Electrophoresis (PFGE) method to purify cellular DNA. From the left: qPCR of the HIV-1 *pol* for the cellular DNA (SupT1 cells) spiked with HIV-1 all-in-one plasmid DNA, circular or linearized. Data shown before and after PFGE as HIV-1 *pol* copies per million *CCR5* copies; representative gel image of the DNA before and after PFGE. Data were statistically tested using a non-parametric Mann–Whitney test p value <0.05 was considered significant, ** = p <0.01. (b) Mapping of large internal deletions (LID) within LTatC[M]L in mCherry+ SupT1 and mCherry+ K562 CRISPRa-NC and -HLTF cells. SupT1 cells were treated with SAHA and TNFα, K562 CRISPRa cells were untreated. Black horizontal bars represent amplified and sequenced regions of the vector, and LID are shown as white horizontal bars. Abbreviations: *cHS4*, chicken hypersensitive site 4; *heIF4A1*, human eukaryotic initiation factor 4A1 gene promoter; *IRES*, internal ribosome entry site; *RRE*, HIV-1 *rev* responsive element; *sMAR8*, synthetic matrix attachment region 8 (from the human β-interferon gene); *tat*, HIV-1 transactivator; *tetO*, tetracycline operator sequences; *WPRE*; woodchuck hepatitis virus posttranscriptional regulatory element. Created with Biorender.

Following purification, DNA from sorted K562-VPR and SupT1 cells was diluted to obtain approximately a single copy of proviral DNA per PCR reaction. We then amplified a region spanning from the 5’ LTR to the start of the mCherry coding sequence, including the entire Cerulean cassette. NGS of the amplified products revealed LID within the target region derived from single mCherry+ cells (Figure 5b), regardless of the treatment with latency-reversing agents, whereas amplicons from double-positive Cerulean+/mCherry+ cells were predominantly intact, as anticipated (data not shown). The LID were over 2,000 bp long, encompassing HIV-1 Tat and Cerulean cassette, similarly to the defects observed in our previous studies employing HIV-1 based dual-fluorophore vectors and in PWH [10–13, 17, 18]. Notably, homologous repeats at the deletion junctions were detectable in only one case and absent in the others, consistent with our hypothesis that cellular DNA damage response proteins, such as HLTF and RAD1, are involved in LID formation.

## Discussion

HIV-1 integration and its interaction with host cellular factors have been studied extensively, and the composition of the latent reservoir, including the high proportion of defective proviruses, is well documented in people with HIV. Internal deletions and other defects in proviral genomes are typically attributed to errors during reverse transcription. However, the potential contribution of host DNA damage response (DDR) and repair pathways to the generation of such defects, particularly large internal deletions (LID) spanning substantial regions of the viral genome, has been relatively underexplored.

In this study, we used an HIV-1-based latency vector, LTatC[M]L, that distinguishes cells harbouring intact vector (double-positive Cerulean+/mCherry+) and defective vector (single mCherry+) via fluorophore signatures to dissect the role of DDR genes in the formation of LID in the HIV-1 provirus. We performed a pooled CRISPR knockout screen targeting DDR genes in SupT1 cells and identified candidate genes that, upon silencing, shifted the balance between single mCherry+ and double-positive cells, indicating that they are potentially involved in the LID formation. Based on reproducibility across three biological replicates, gene function, and reported or hypothesised interactions with HIV-1, we selected 10 candidate genes for follow-up. We then functionally validated these candidates using both CRISPR activation and siRNA-mediated knockdown. In the CRISPRa experiments, overexpression of *HLTF*, *TREX2*, *RAD18*, and *PSIP1 (LEDGF)* consistently increased the proportion of single mCherry+ cells, indicating an increased frequency of defective proviruses, with *HLTF* showing the strongest and most statistically robust effect. Several other genes showed trends in the same direction but did not reach significance, in part due to variability across replicates. *BARD1* and *RAD1* could not be reliably overexpressed, limiting the interpretation of their CRISPRa phenotypes. Complementary siRNA experiments reinforced *HLTF* as a key candidate and additionally supported a role for *RAD1*, whose knockdown significantly affected the distribution of defective versus intact proviruses, particularly in combination with *HLTF*. Although effect sizes were modest, they were reproducible, which is consistent with DNA repair proteins being tightly regulated and partially buffered by redundancy.

Our CRISPRa data suggest that increased *PSIP1 (LEDGF)* expression modestly enhances the formation or detection of defective proviruses, whereas siRNA-mediated *PSIP1 (LEDGF)* depletion had no significant effect on the ratio of defective (single mCherry+ cells) to intact (double-positive Cerulean+/mCherry+ cells) proviruses. PSIP1 (LEDGF) promotes efficient HIV-1 integration into actively transcribed genes [29], and integration is reduced but not abolished upon PSIP1 (LEDGF) loss, with partial compensation by HRP-2 [38]. One possible explanation is that *PSIP1 (LEDGF)* overexpression increases integration in PSIP1 (LEDGF)-preferred, transcriptionally active loci and thereby provides more substrates for DNA repair-linked processing that generates LID, in line with our hypothesis that defects arise during or shortly after integration. Conversely, when *PSIP1 (LEDGF)* is silenced, overall integration is reduced and a greater fraction of residual integration may occur at transcriptionally weaker or PSIP1 (LEDGF)-independent sites (for example, near promoters or CpG islands), leading to proviruses that are poorly expressed and thus fall below our fluorophore-based detection, so the observed single mCherry+ to double-positive cell ratio remains essentially unchanged despite altered integration patterns.

HLTF stood out as the most consistent hit across our perturbation strategies: CRISPR activation of *HLTF* increased the proportion of defective (single mCherry+) proviruses, whereas siRNA-mediated knockdown reduced this population. HLTF is a member of the human SWI/SNF (SWItch/Sucrose Non-Fermentable) protein family involved in ATP-dependent chromatin remodelling. Multiple roles of HLTF have been described, including promoting DNA damage tolerance via replication fork reversal, and transcriptional modulation, with the latter function previously linked to its binding of the HIV-1 promoter [39–42].

HLTF is increasingly recognised as a broad antiviral restriction factor, acting through DNA repair-linked mechanisms in HIV-1 and HCMV and through modulation of autophagy in HTLV-1, and is actively antagonized by viral proteins in each case [27, 43]. We do not propose HLTF as an alternative to the classical model in which defects originate from errors during reverse transcription; instead, our data support an expanded view in which host DDR proteins, including HLTF, act on viral DNA in addition to the intrinsic error-prone nature of reverse transcription. In this view, viral reverse transcription and integration generate gapped, nicked, and branched DNA intermediates that are then further processed by host factors, and in a fraction of events, this results in LID formation rather than faithful repair.

On the basis of published work on HLTF recognition of fork-like or gapped DNA [27] and our own phenotypes, we consider two non-mutually exclusive mechanistic scenarios. In model 1, HLTF acts on late reverse transcription intermediates before integration: branched, flap-containing, and gapped viral DNA structures resemble stalled replication forks, and HLTF, via its HIRAN domain and helicase activity, can bind and remodel such intermediates. Excessive binding and unwinding in the context of *HLTF* overexpression could expose longer stretches of single-stranded or misaligned viral DNA to nucleases, leading to trimming of internal regions; the resulting shortened molecules can still integrate but manifest as large internal deletions in single mCherry+ cells. In model 2, HLTF acts shortly after integration on gapped proviral DNA encountering a host replication fork: early proviruses may initially be incompletely repaired, and an advancing host fork colliding with a not-yet fully double-stranded proviral DNA could recruit HLTF to stabilize and remodel the stalled fork. In most cases, this would promote repair, but in a subset of events, fork reversal and error-prone resection or template switching could remove internal segments of viral DNA, again generating LID; overexpression of *HLTF* would increase such events, whereas depletion would reduce them.

Other factors identified in our screen fit naturally into this framework, such as RAD1, TREX2, ZRANB3 (belongs to the SNF2 family, same as HLTF), and RAD18, whether they reach statistical significance or not. They support a model in which multiple DNA damage response pathways cooperate with HLTF to process viral DNA intermediates, with large deletions arising from the combined action of several host enzymes on structurally abnormal viral DNA rather than from a single polymerase error.

Sequencing of double-positive Cerulean+/mCherry+ and single mCherry+ populations confirmed that they predominantly contain intact and internally deleted proviral genomes, respectively, consistent with our previous work [17, 18]. Deletions frequently remove large portions of the Cerulean cassette and can extend from the 5’ LTR into the mCherry region, and short direct repeats at deletion junctions, expected from classical strand slippage during reverse transcription, were rare, again suggesting that additional mechanisms beyond reverse transcription errors contribute to these defects [17, 18].

Our study relies on a lentiviral vector system in immortalized cell lines, and we selectively analyse integrated, transcriptionally detectable proviruses. These choices allowed us to dissect the mechanism of LID formation in a controlled setting, but future work in primary cells and patient-derived samples will be essential to define how far the same DNA repair pathways shape defective proviruses *in vivo*. It will also be important to move more systematically from single-gene to combinatorial perturbations, testing how simultaneous modulation of HLTF with other repair factors affects the LID frequency in both vector and replication-competent HIV-1. A particularly attractive next step is to dissect HLTF’s mechanism in more detail by combining knockout and rescue approaches with catalytically impaired or domain-specific *HLTF* mutants, and by introducing HIV-1 Vpr into our system to test directly how viral counteraction of HLTF influences the generation of LID; ideally, these experiments would be coupled to quantitative analyses of reverse-transcription and integration intermediates to distinguish pre- versus post-integration modes of action.

Despite modest effect sizes and the constraints of our model system, our data highlight an underexplored aspect of HIV biology: host DNA repair and damage-response factors can act as restriction mechanisms that favour the accumulation of defective proviruses rather than simply repairing viral DNA. We view this work as a starting point, pointing toward a broader role of cellular repair pathways in sculpting the proviral landscape and raising the possibility that these pathways could be modulated therapeutically to bias infection outcomes toward non-productive, defective integrations.

## Materials & Methods

### Plasmid preparation

LTatC[M]L was cloned as described previously [18]. LentiGuide-Puro-P2A-EGFP was a gift from Fredrik Wermeling (Addgene plasmid #137729, [44]), psPAX2 from Didier Trono (Addgene plasmid #12260), and pCAG-VSVG from Arthur Nienhuis and Patrick Salmon (Addgene plasmid #35616). Single mCherry and Cerulean plasmid constructs in the LTatCL[M] backbone, used for FACS compensation, were generated by excising one of the fluorophores, accordingly. In the single Cerulean construct, the HIV-1 5’ LTR promoter was replaced with the human promoter, eF1α. HIV-1 all-in-one standard plasmid was generated by Dr. Victoria Strouvelle [37].

CRISPRko DNA damage response library (DDR, DNA Damage Response MKOv4 Library) was a gift from Junjie Chen (Addgene pooled library #140219). The library was amplified according to the Broad Institute protocol (Amplification of pDNA libraries). Briefly, 400 ng of library DNA was added to 100 μl of STBL4 electrocompetent cells (ElectroMAX™ Stbl4™ Competent Cells, Thermo Fisher Scientific, 11635018), electroporated using 1.8 kV program (MicroPulser^TM^, Bio-Rad), recovered in 10 ml of SOC for 1 h at 30°C, and plated. Following overnight incubation at 37°C, bacterial colonies were scraped with cold LB broth (Sigma-Aldrich, L3022) and directly used for DNA isolation by QIAGEN Plasmid Maxi Kit (Qiagen, 12165).

### Cell culture

SupT1 cells, ([45] kindly provided by Dharam Ablashi through the NIH AIDS Reagent Programme, Division of AIDS, NIAID, NIH) and K562 cells (American Type Culture Collection, CCL-243 ™) were cultured in RPMI-1640 (Sigma-Aldrich, R8758) supplemented with 10% (v/v) fetal bovine serum (FBS) (Thermo Fisher Scientific, A5256701), 100 U/ml Penicillin (Gibco™, 15140122), and 100 μg/ml Streptomycin (Gibco™, 15140122). K562-VPR cells (Horizon Discovery, HD dCas9-VPR-005) were cultured in IMDM (Gibco™, 12440053) supplemented with 10% (v/v) FBS, 100 U/ml Penicillin, and 100 μg/ml Streptomycin. 293T cells (American Type Culture Collection, CRL-3216™) were cultured in DMEM (Sigma Aldrich, D6429) supplemented with 10% (v/v) FBS, 100 U/mL Penicillin, and 100 μg/ml Streptomycin. All cell lines were kept in culture at 37°C, 5% CO2, and split every 2-3 days in a ratio of 1:10

### Virus production and titration

VSV-G-pseudotyped LTatC[M]L, CRISPRko DDR library, and CRISPRa-gRNA (LentiGuide-Puro-P2A-EGFP) lentiviral vectors were produced by transfection of 293T cells. On the day of transfection, 293T cells were at least 80% confluent in T-150 flasks and were transfected with 20.8 μg transfer plasmid (LTatC[M]L, CRISPRko DDR library, or CRISPRa-gRNA, respectively), 20.8 μg psPAX2, 10.4 μg pCAG-VSVG, and 208 μg polyethylenimine (PEI 25K; Polysciences, MW 25,000, transfection grade) in 3 ml serum-free DMEM. The media was replaced with complete DMEM approximately 18 h post-transfection. Virus-containing supernatants were harvested on days 3 and 4 post-transfection, passed through a 0.22 μm filter, and concentrated by incubation with 1x PEG-it (System Biosciences, LV825A-1-SBI) at 4°C overnight. Viral particles were pelleted by centrifugation at 1,500 × g for 1 h at 4°C and resuspended in 450 μl RPMI-1640.

CRISPRko DDR library virus stocks were titrated by transducing 40,000 SupT1 cells in 50 μl media supplemented with 8 μg/ml polybrene (Sigma-Aldrich, TR-1003-G) in 96-well plates, using serial dilutions of viral stock up to 1:10,000 in quadruplicate. Cells were cultured with or without 6 μg/ml puromycin (Sigma-Aldrich, P7255), and viable cells were counted on day 5 post-transduction. The percentage of transduced cells was inferred by dividing the number of viable cells in puromycin-treated wells by that in untreated wells.

LTatC[M]L virus stocks were titrated by transducing 0.5 × 10^6^ K562 or K562 VPR cells in 500 μl media supplemented with 8 μg/ml polybrene in 12-well plates, using serial dilutions of viral stock up to 1:100,000 in triplicate. CRISPRa-gRNA (LentiGuide- Puro-P2A-EGFP) virus stocks were titrated by transducing 0.25 × 10^6^ K562-VPR cells in 500 μL media supplemented with 8 μg/ml polybrene in 24-well plates, using serial dilutions of viral stock up to 1:1,000 in triplicate. On day 5 post-transduction, the percentage of fluorescent cells was quantified by flow cytometry (CytoFLEX S, Beckman Coulter): mCherry and/or Cerulean positivity for LTatC[M]L and GFP positivity for CRISPRa-gRNA. For viral titre calculations, only wells with <20% positive cells were used to ensure a high probability of single integration events per cell [46]. Viral titres were expressed as transduction units per millilitre (TU/ml) and were used for MOI calculation.

### CRISPR/Cas9 screening

#### CRISPR/Cas9 pooled screening

A total of 1.5 × 10^7^ SupT1 cells were transduced with VSV-G-pseudotyped CRISPRko DDR library, as described by Su et al., at an MOI of 0.3 in 20 mL media supplemented with 8 μg/ml polybrene [19]. Following overnight incubation, the cell culture media was supplemented with 2 μg/ml puromycin. After 72 h of puromycin selection, a minimum 1.5 × 10^7^ cells were transduced with VSV-G-pseudotyped LTatC[M]L at an MOI of 0.5 in 20 ml media. At least 3 × 10^6^ cells were collected at this stage as a background control sample. On day 10 post-LTatC[M]L transduction, double-positive Cerulean+/mCherry+ cells and single mCherry+ cells were bulk-sorted by flow cytometry (BD FACSymphony S6). Genomic DNA was extracted directly from all samples, including the background control, using the DNeasy Blood & Tissue Kit (QIAGEN, 69504), yielding a minimum of 4.5 μg DNA per sample. For the third biological replicate (DDR 3), an additional sorting step was performed after CRISPRko DDR library transduction to remove dead cells based on forward and side scatter (BD FACSymphony S6).

#### Sequencing

Genomic DNA was quantified using a NanoDrop One C microvolume spectrophotometer (Thermo Scientific). To achieve the desired library coverage, 9 μg gDNA per sample was amplified in parallel one-step PCRs with barcoded primers targeting the lentiCRISPRv2 vector backbone (primer sequences in Table 1). Primers were ordered as desalted oligonucleotides from IDT (Integrated DNA Technologies, LubioScience GmbH). Each 100 μL PCR contained 1.8 μg gDNA and 0.02 U/μl Q5 polymerase (New England Biolabs) and was run for 23–27 cycles.

**Table 1.**
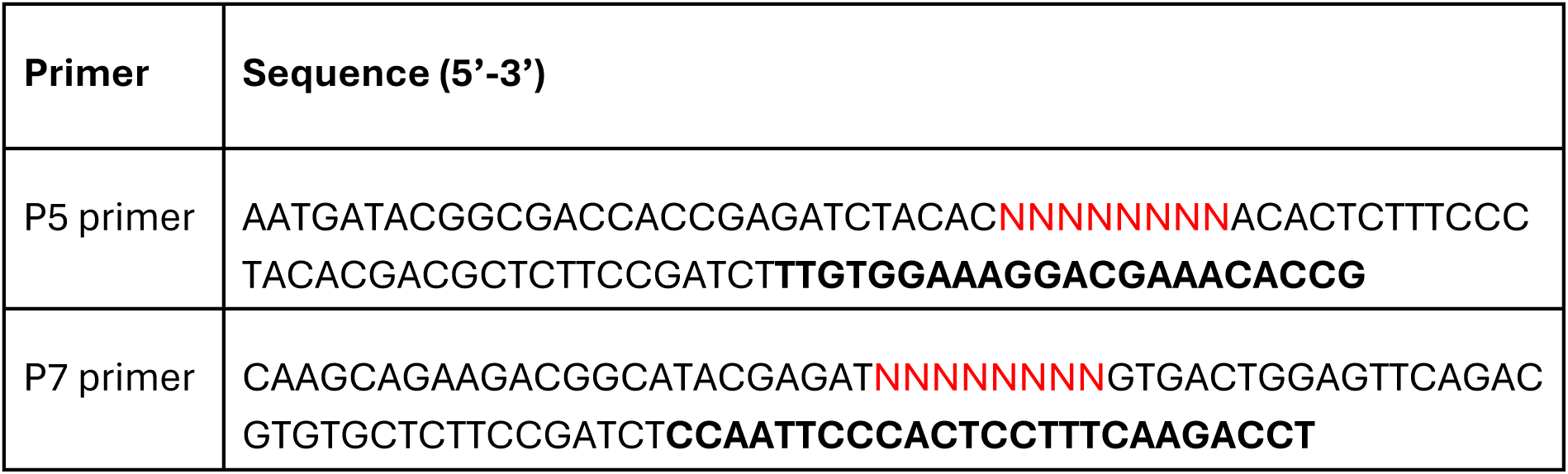
Primer sequences used for PCR amplification. 8 bp barcode position is marked in red, vector binding site in bold.

PCR products containing the spacer sequences were purified by SPRI bead cleanup with a right-sided size selection to remove genomic DNA input. Using a dual-index barcoding strategy, NGS amplicons were multiplexed and sequenced on an Illumina platform with 21 dark cycles (DC) in Read 1 to skip the invariable U6 promoter region. Sequencing parameters were 21 DC + 36 bases for Read 1, 8 bases for Index 1 (I1), and 8 bases for Index 2 (I2), with a 5–10% PhiX Control v3 spike-in (Illumina). Libraries were loaded on a NextSeq 2000 at 350 pM. BCL files were demultiplexed using bcl2fastq (Illumina), allowing 0 mismatches in Index 1 and 1 mismatch in Index 2. Data quality was evaluated with FastQC (Babraham Bioinformatics) to assess base quality scores and other standard QC metrics. In addition, an in-house pipeline mapped a subset of sgRNA reads against commonly used CRISPR libraries using the R package ezRun (https://github.com/uzh/ezRun) within the SUSHI data analysis framework [47].

#### Data processing

Next-generation sequencing reads were aligned to the CRISPRko DDR library, and sgRNA counts were generated. Data quality control, single-gRNA counting, and gene-level statistical testing and scoring were performed using the MAGeCK algorithm [20].

### Cloning of CRISPR activation guide RNAs

For each target gene, three guide RNAs (gRNAs) were selected from commercially available CRISPR activation libraries (Calabrese and SAM) [33, 34], and one non-targeting gRNA was designed as a negative control. gRNA sequences were evaluated using the CHOPCHOP web tool (https://chopchop.cbu.uib.no/) to minimize predicted off-target sites and self-complementarity and to maximize on-target efficiency. The final gRNA sequences (without added overhangs) and corresponding forward and reverse oligonucleotides are listed in Supplementary Table S5.

gRNAs were cloned separately into the LentiGuide-Puro-P2A-EGFP backbone (Addgene 137729) using Golden Gate cloning. The plasmid was digested with BsmBI-v2 (New England Biolabs, R0739S), which cuts between the U6 promoter and the gRNA scaffold. To generate compatible overhangs, the forward oligonucleotide consisted of the gRNA sequence preceded by “CACCG”, where “CACC” provides the appropriate overhang, and the additional “G” supports transcription initiation from the U6 (Pol III) promoter. The reverse oligonucleotide was the reverse complement of the gRNA sequence with “AAAC” appended at the 5’ end, matching the overhang created on the scaffold side, and an additional “C” at the 3’ end to pair with the initiating “G” in the forward strand. Oligonucleotides were ordered from Microsynth AG and dissolved in Tris buffer to a concentration of 100 µM.

Single gRNAs were cloned into LentiGuide-Puro-P2A-EGFP following the Zhang lab SAM cloning protocol provided by Addgene (Zhang lab SAM cloning protocol, Addgene, [48]), with minor modifications to reagent sources and vector backbone. The Golden Gate reaction contained 25 ng backbone plasmid, 1 µl of 1:10 diluted annealed gRNA oligonucleotides, 0.5 µl BsmBI-v2, 0.5 µl T4 DNA ligase (Thermo Scientific, EL0011), 1 µl 10 mM ATP (New England Biolabs, P0756S), 1 µl NEB Buffer r3.1 (New England Biolabs), and nuclease-free water to the final reaction volume.

A volume of 1 µl of the Golden Gate reaction was transformed into Stellar Competent Cells (TaKaRa Bio, 636766) according to the manufacturer’s Stellar Competent Cells protocol (PT5055-2). Transformed cells were plated on LB agar plates containing 100 µg/ml ampicillin (ampicillin sodium salt, Sigma-Aldrich, A0166) and incubated overnight at 37°C. The following day, one to two colonies per construct were inoculated into LB broth (Sigma-Aldrich, L3022) supplemented with ampicillin and grown overnight. Plasmid DNA was isolated using the QIAGEN Plasmid Maxi Kit (Qiagen, 12165) according to the manufacturer’s instructions. Plasmid DNA concentration was measured by NanoDrop and Qubit (Qubit 4, Invitrogen), and correct gRNA insertion was confirmed by NGS.

### siRNA

K562 cells were seeded on the day of transfection at 1 × 10^5^ cells per well of a 48-well plate in 50 μl media. The cells were transfected with 3 μl of gene-targeting siRNA (Santa Cruz Biotechnology) or non-targeting control siRNA (Santa Cruz Biotechnology, control siRNA-A, sc-37007), and 4.5 μl HiPerFect Transfection Reagent (QIAGEN, 301704), which were diluted in 50 μl of Opti-MEM^TM^ Reduced Serum Medium (Gibco^TM^, 31985062). The media was changed after 5 h of incubation. The following gene-targeting siRNAs were ordered from Santa Cruz Biotechnology, with each being a pool of three siRNA duplexes: sc-44991 (*PSIP1*), sc-72142 (*RAD18*), sc-94423 (*ZRANB3*), sc-36356 (*RAD1*), sc-45943 (*HLTF*).

The cells were transduced with LTatC[M]L between 29 h and 51 h post-siRNA transfection. For functional assays, 4 x10^4^ cells from each siRNA-transfected population were transduced with LTatC[M]L lentivirus at an MOI of 0.5 in 50 μl cell culture media supplemented with 8 μg/ml polybrene in 96-well plates, in triplicate. Cells were expanded and analysed on day 7 post-transduction for double-positive Cerulean+/mCherry+ and single mCherry+ expression. Minimum three biological replicates were generated and analysed.

### CRISPR activation

For each candidate gene, three gRNAs were cloned individually into the CRISPRa expression vector LentiGuide-Puro-P2A-EGFP, pooled, and packaged into lentiviral particles. 5 × 10^6^ K562-VPR cells were transduced at an MOI of 0.5, and GFP+ cells were isolated by FACS to obtain K562-VPR populations overexpressing a single candidate gene. Gene and protein expression of cells were measured. For functional assays, 4 × 10^4^ cells from each GFP+ population were transduced with LTatC[M]L lentivirus at an MOI of 0.5 in 50 μl media supplemented with 8 μg/ml polybrene in 96-well plates, in triplicate. Cells were expanded and analysed on day 7 post-transduction for double-positive Cerulean+/mCherry+ and single mCherry+ expression, and changes in the ratio of these populations relative to cells transduced with non-targeting gRNAs were quantified and statistically analysed. Three biological replicates were generated and analysed.

### Flow cytometry and cell sorting

Cells transduced with lentiviral vectors carrying either the model vector LTatC[M]L or the GFP-expressing CRISPRa-gRNA backbone (lentiGuide-Puro-P2A-EGFP) were analysed on a CytoFLEX S flow cytometer (Beckman Coulter) to measure GFP, mCherry, and Cerulean fluorescence. When K562-VPR cells were sequentially transduced with the GFP gRNA vector and LTatC[M]L, spectral overlap between GFP and Cerulean was minimized by compensation using single-colour controls. For the Cerulean and mCherry controls, cells were nucleofected with Cerulean-only and mCherry-only plasmids using an Amaxa Nucleofector II (Amaxa Biosystems, Lonza) and Nucleofector 2b Kit V (Lonza, VCA-1003) with program T-016 according to the manufacturer’s protocol for K562 cells. For the GFP single-colour control, cells transduced only with the GFP CRISPRa-gRNA plasmid were used. A compensation matrix was generated in CytExpert software (Beckman Coulter) and applied to all samples prior to data acquisition. Gating strategies were kept constant across experiments.

GFP-positive K562-VPR cells (indicating successful integration of gRNA vectors) were isolated by fluorescence-activated cell sorting (FACS) at the Cytometry Facility, University of Zurich, using a BD FACSAria III (BD Biosciences) at University Hospital Zurich. Similarly, single mCherry+ and double-positive Cerulean+/mCherry+ populations were sorted from 5 × 10^6^ GFP+ K562-VPR and SupT1 cells transduced with LTatC[M]L at an MOI of 0.5 for downstream analysis. SupT1 cells were additionally reactivated with 10 ng/ml TNFα (tumour necrosis factor) and 1 µM SAHA (Suberoylanilide hydroxamic acid) prior to sorting. Given the minimal spillover observed between GFP and Cerulean channels under the chosen configuration, sorting on the BD FACSAria III was performed without additional compensation. All flow cytometry data, including gating, were analysed using FlowJo v10.0.08 (BD Biosciences).

### RT-qPCR

#### RNA isolation and reverse transcription to cDNA

Total RNA was isolated from 5 × 10^5^ (untreated and siRNA-treated samples) to 3 × 10^6^ cells (CRISPRa-treated samples) using the RNeasy Mini Kit (Qiagen, 74104) according to the manufacturer’s protocol, with two on-column DNase treatments (DNase I recombinant, RNase-free, Roche, 04716728001). PBS-only samples were processed in parallel as negative controls. After cell lysis and homogenization, the lysate was mixed with DNase incubation buffer (39 µl 10x DNase buffer and 3 µl DNase I) and incubated at room temperature for 45 min. The sample was then applied to a RNeasy Mini spin column, and RNA purification was continued according to the standard Qiagen protocol. A second DNase treatment was performed after the first RW1 wash. Instead of loading the full 700 µl Buffer RW1, only 350 µl was added, and the column was centrifuged at 8,000 × g for 15 s. Then, 80 µl DNase mix (69 µl RNase-free water, 8 µl 10x DNase buffer, 3 µl DNase I) was applied directly to the column membrane and incubated at room temperature for 45 min. The remaining 350 µl of Buffer RW1 were then added, the column was centrifuged, and the protocol was continued with Buffer RPE washes as recommended by Qiagen. RNA was eluted in 25 µl RNase-free water.

cDNA was synthesized using PrimeScript Reverse Transcriptase (TaKaRa Bio, 2680B) and random hexamer primers (hexadeoxyribonucleotide mix, pd(N)6; TaKaRa Bio, 3801). No-RT controls (lacking reverse transcriptase) were processed in parallel. Each reaction contained 4 µl 5x PrimeScript Buffer, 1 µl 10 mM dNTP mix (dNTP Set, 100 mM solutions, Thermo Scientific, R0182), 1 µl random primers (100 µM), and 0.2 µl PrimeScript RTase (omitted in no-RT controls). The volume was adjusted to 15 µl with RNase-free water, and 5 µl RNA was added to each reaction, including PBS controls. Reverse transcription was carried out in a thermal cycler under the following conditions: 37°C for 15 min, 85°C for 5 s, then hold at 4°C.

#### qPCR

To assess mRNA expression of overexpressed or silenced genes or baseline expression of candidate genes in SupT1 and K562 cells, quantitative PCR (qPCR) was performed using SYBR Green chemistry (Thermo Scientific) and gene-specific standards on a QuantStudio 5 or ABI 7500 Real-Time PCR System (Applied Biosystems). Gene-specific primers (Microsynth AG) were designed spanning exon-exon junctions to amplify a defined region of each target, including the housekeeping gene *β-actin*; primer sequences and cycling conditions are listed in Supplementary Table S6.1. Amplicons were visualized on agarose gels, purified using the NucleoSpin Gel and PCR Clean-up kit (Macherey-Nagel, 740609.50), and verified by Sanger sequencing (Microsynth AG) or NGS, as appropriate. These purified, sequence-confirmed amplicons were used to generate standard curves (10^7^ to 10^1^ molecules/µl) for absolute quantification of gene expression.

Each qPCR reaction contained 5 µl 4x Real-Time Premix (including 0.02 µM ROX [6-ROX, Invitrogen, C-6156], 4x PCR buffer [10x PCR buffer without MgCl₂, Sigma-Aldrich, P2317], 6 mM MgCl₂ [25 mM MgCl₂ solution, Sigma-Aldrich, M8787], 1.6 mM dNTP mix [Thermo Fisher Scientific, R0181]), 0.4 µl 10x SYBR Green (SYBR Green I, 10,000x, Invitrogen, S-7585), 0.4 µl each of 20 µM forward and reverse primers (Microsynth AG), and 0.2 µl JumpStart Taq DNA polymerase (Sigma-Aldrich, D6558), adjusted to 10 µl with nuclease-free water. To each reaction, 10 µl of either 1:10 diluted cDNA or a standard dilution was added. No-RT and PBS controls were included in every run. All samples, standards, and controls were run in duplicate. Primer sequences, amplicon lengths, and qPCR cycling conditions are provided in Supplementary Table S6.2. Data were analysed using QuantStudio Design and Analysis Software or ABI 7500 Real-Time PCR Software v2.3. Gene expression was normalized to housekeeping genes (*GAPDH* or *β-actin*) and compared between samples and non-targeting gRNA controls to determine fold changes in expression. For *GAPDH* quantification, purified amplicons of the HIV-1 all-in-one standard were used to generate the corresponding standard curve.

qPCR of the HIV-1 *pol* gene was also performed using TaqMan chemistry on a QuantStudio 5 Real-Time PCR System (Applied Biosystems) to assess the efficiency of BluePippin size selection. Serial dilutions of the purified amplicon of HIV-1 all-in-one standard plasmid were used to generate qPCR standards. Primers, probes, reagent concentrations, and cycling conditions are listed in Supplementary Table S6.3. qPCR data were normalized to copies per 10^6^ *CCR5* copies and compared between samples before and after PFGE.

### Western blot

Cells (1 × 10^6^ per well) were seeded in 6-well plates and incubated at 37°C, 5% CO₂. Cells were harvested the next day, centrifuged at 300 × g for 5 min, and the supernatant was discarded. Pellets were washed once with PBS, centrifuged again, and the cell pellet was collected. A total of 150 µl RIPA buffer (Pierce RIPA Lysis and Extraction Buffer, Thermo Scientific, 8990) supplemented with 1.5 µl of 100x Halt protease and phosphatase inhibitor cocktail (Thermo Scientific, 78440; diluted 1:100 in RIPA) was added. Samples were vortexed and incubated on ice for 15 min with intermittent vortexing. Lysates were centrifuged at 16,000 × g for 20 min at 4°C, and the supernatant was collected. Protein concentration was determined using the Pierce BCA Protein Assay Kit (Thermo Scientific, 23227).

For SDS-PAGE, 15 µg (CRISPRa samples) or 20 µg (siRNA-treated samples) of protein was mixed with 4x Bolt LDS Sample Buffer (Thermo Scientific, B0007) supplemented with 50 mM DTT to a final volume of 30 µl, yielding a final 1x LDS concentration. Samples were incubated at 70-80°C for 10 min. Proteins were resolved on Bolt 4-12% Bis-Tris Plus Mini Protein Gels (15- well, Invitrogen, NW04125BOX) using an XCell SureLock Mini-Cell (Thermo Scientific, EI0001). Bolt MES SDS running buffer (20x, diluted to 1x with distilled water; Invitrogen, B0002) was used as running buffer, and 5 µl AcuteBand Pre-Stained Protein Ladder (Lubio Science, LU5001-0500) was loaded as a molecular weight marker. A total of 30 µl sample per lane was loaded, and gels were run for 20 min at 80 V, followed by 60 min at 120 V.

Proteins were transferred to nitrocellulose membranes (Transfer Membrane ROTI® NC 0.45, 300 × 30 cm, Carl Roth, 200K.1) by wet electroblotting using 10% methanol in 1x Bolt Transfer buffer (20x, diluted with distilled water; Invitrogen, BT00061) and an XCell II Blot Module (Invitrogen, EI9051) at 30 V for 1 h (current limit 200 mA). After transfer, membranes were blocked in 5% BSA in 1 × TBS-0.1% Tween-20 (TBS-T) (10x TBS, Canvax, BR0033; Tween-20, Sigma-Aldrich, P1379) for ≥1 h at room temperature or overnight at 4°C.

Blocked membranes were incubated with primary antibodies diluted in 5% BSA in 1x TBS-T on a shaker, typically overnight at 4°C. The next day, membranes were washed in 1x TBS-T (3 × 5min), incubated with HRP-conjugated secondary antibodies diluted in 5% BSA in 1x TBS-T for ≥1 h at room temperature, and washed again in 1x TBS-T (3 × 5min). Antibody dilutions and incubation conditions followed the manufacturer’s datasheets.

Signals were developed using Pierce ECL Western Blotting Substrate (Thermo Scientific, 32209) and imaged on a Vilber Fusion FX7 chemiluminescence imager. Data were analysed using EvolutionCapt software. Target protein expression was normalized to α-tubulin and compared between samples and non-targeting controls to calculate fold changes. In some experiments, membranes were treated with Restore PLUS Western Blot Stripping Buffer (Thermo Scientific, 46430) according to the manufacturer’s instructions before incubation with α-tubulin antibody, to remove target proteins such as PSIP1 (LEDGF) and RAD18 that migrate at a similar apparent molecular weight to α-tubulin.

Primary antibodies used were: HLTF (G-6, mouse monoclonal IgG1κ, Santa Cruz Biotechnology, sc-398357; 1:100), PSIP1 (3F7, mouse monoclonal IgG1κ, Santa Cruz Biotechnology, sc-101087; 1:200), RAD18 (79B1048, mouse monoclonal IgG1, Santa Cruz Biotechnology, sc-52949; 1:200), ZRANB3 (rabbit polyclonal IgG, Elabscience Bionovation, E-AB-65632; 1:2000), TREX2 (E-4, mouse monoclonal IgG1κ, Santa Cruz Biotechnology, sc-390890; 1:100), RAD1 (D-6, mouse monoclonal IgG2aκ, Santa Cruz Biotechnology, sc-166515; 1:100), and α-tubulin (DM1A, mouse monoclonal IgG1κ, Abcam, ab7291; 1:10,000). Secondary antibodies used were HRP-linked horse anti-mouse IgG (Cell Signaling Technology, 7076P2; 1:1000) and HRP-linked goat anti-rabbit IgG (Cell Signaling Technology, 7074S; 1:3000).

### Pulsed-field gel electrophoresis

Pulsed-field gel electrophoresis (PFGE) was used to enrich for high-molecular weight cellular DNA containing integrated copies of the model vector and to remove episomal or unintegrated DNA fragments. PFGE was performed using a BluePippin instrument (Sage Science). Reagents (electrophoresis buffer, elution buffer, loading solution, and DNA marker) and gel cassettes were obtained from Sage Science (kit PN/catalog number BLF7510). The protocol was followed according to the manufacturer’s instructions, using the cassette definition “0.75% DF Marker S1 high-pass 15-20 kb” to collect DNA >15 kb and deplete shorter or sheared DNA fragments.

Genomic DNA was isolated using the DNeasy Blood and Tissue Kit (Qiagen, 69506), quantified by NanoDrop, concentrated by evaporation, and re-quantified using a Qubit fluorometer (Qubit 4, Invitrogen). Between 500 ng and 5 µg DNA (depending on availability) in a maximum volume of 30 µl was loaded per lane for PFGE, in accordance with the instrument specifications (maximum 5 µg per lane). Eluted DNA fractions were collected and quantified again using Qubit.

To verify the efficiency of the PFGE size-selection step, SupT1 cellular DNA was spiked with linearized or circular forms of a laboratory-designed “HIV-1 all-in-one standard” plasmid containing *GAPDH*, *CCR5*, *LTR*, *pol*, and *gag* sequences. Samples taken before and after BluePippin processing were analysed by agarose gel electrophoresis and quantified for the HIV-1 *pol* region by qPCR to assess removal of shorter DNA fragments and plasmid DNA, using three independent biological replicates for each plasmid condition.

### Mapping of large internal deletions

PFGE-purified DNA from double-positive Cerulean+/mCherry+ and single mCherry+ cells was diluted to achieve approximately a single vector copy per reaction and amplified by PCR to map deletions within the Cerulean cassette. For each DNA sample, at least 16 independent PCR reactions were performed. Each 15 μl PCR contained 5 μl 5x LongAmp Taq Reaction Buffer (New England Biolabs, M0323L), 0.75 μl 10 mM dNTP mix (Thermo Fisher Scientific, R0181), 1.25 μl 10 μM forward primer, 1.25 μl 10 μM reverse primer, and 1 μl LongAmp Taq DNA Polymerase (NEB, M0323L), and was brought to volume with nuclease-free water; 10 μl of diluted DNA was added per reaction.

PCR amplicons were resolved by agarose gel electrophoresis to assess product size, then purified using the NucleoSpin Gel and PCR Clean-up Kit (Macherey-Nagel, 740609.50) and submitted for NGS. Primer sequences and cycling conditions are listed in Supplementary Table S6.4.

### Next-generation sequencing (NGS)

To verify plasmid constructs containing cloned gRNAs and to map deletions in the target region of the model vector, NGS was performed on (i) plasmids generated by Golden Gate cloning and (ii) PFGE-purified DNA from double-positive Cerulean+/mCherry+ and single mCherry+ cells. DNA concentrations were measured with a Qubit fluorometer, and samples were diluted to 0.2ng/µl.

Libraries were prepared using the Nextera XT DNA Library Prep Kit (Illumina, FC-131-1096) and the Nextera XT Index Kit v2 Set A (Illumina, FC-131-2001). Sequencing was performed on an Illumina MiSeq platform using the MiSeq Reagent Kit v3, 150-cycle format (Illumina, MS-102-3001), according to the manufacturer’s specifications and with a 1% PhiX control spike-in.

All plasmid constructs used in this study, including gRNA expression vectors and lentiviral packaging/expression plasmids, were sequence-verified by NGS as described above. Sequencing data were retrieved from the openBIS system (University of Zurich) and analysed using CLC Genomics Workbench 25 (Qiagen).

### Statistics

Gene-level statistical testing and scoring of CRISPR screening were performed using the MAGeCK algorithm [20]. The differences between control gRNA counts in the samples were assessed using a two-tailed non-parametric Mann-Whitney U-Test. The reduction in the HIV-1 pol gene copy number following PFGE was tested using a non-parametric Mann–Whitney test comparing pre- and post-PFGE samples in GraphPad Prism 10. Statistical significance was assessed using p values (α = 0.05). To assess the effect of gene overexpression and silencing, mixed-effects logistic regression models were fitted in R, with the log-odds of being single mCherry+ versus double-positive Cerulean+/mCherry+ as the response, gene as a fixed effect, and biological replicate as a random effect. Statistical significance was assessed using p values (α = 0.05), and multiple-testing correction was performed using the Benjamini–Hochberg procedure; adjusted p values ≤ 0.1 were considered significant.

## Data availability

All data underlying the results presented in this study are included in the article and its Supporting Information files. Raw next-generation sequencing data will be deposited in the European Nucleotide Archive (ENA) and made publicly available upon publication; accession numbers will be provided in the final version of the article. Additional underlying data can be obtained from the corresponding author upon reasonable request.

## Abbreviations

AIDS: Acquired immunodeficiency syndrome
ART: Antiretroviral therapy
CEG: Core-essential genes
CRISPRa: CRISPR activation
CRISPRko: CRISPR knockout
DDR: DNA damage response
gRNA: guide RNA
HIV-1: Human immunodeficiency virus type 1
LID: Large internal deletions
LTatC[M]L: 5’ LTR Tat-Cerulean- insulator-mCherry-insulator-3’ LTR
LTR: Long terminal repeat
MAGeCK: Model-based Analysis of Genome-wide CRISPR/Cas9 Knockout
MOI: Multiplicity of infection
NEG: Non-essential genes
NGS: Next-generation sequencing
NHEJ: Non-homologous end joining
PCA: Principal component analysis
PFGE: Pulsed-field gel electrophoresis
PWH: People with HIV
TNFα: Tumour necrosis factor
SAHA: Suberoylanilide hydroxamic acid
siRNA: Small interfering RNA

## Acknowledgements

We thank Philipp Schätzle, Manuel Schulthess, Mario Wickert, Tatiane Gorski (Flow Cytometry Facility, UZH, Zurich, Switzerland) for bulk cell sorting services, Gabriela Ziltener (Institute of Medical Virology, UZH, Zurich, Switzerland) for next-generation sequencing, Aria Maya Minder Pfyl (Genetic Diversity Centre, ETH, Zurich, Switzerland) for the use of the BluePippin device, and Tom Loosli for sharing R script used for the gRNA diversity analysis.

The following reagent was obtained through the NIH AIDS Reagent Programme, Division of AIDS, NIAID, NIH: SupT1 from Dr. Dharam Ablashi.

## Author Contributions

- Koleta Michalek conceived and designed the experiments, performed the experiments, analysed the data, prepared figures and/or tables, authored or reviewed drafts of the paper, and approved the final draft.
- Sagnik Bhattacharjee conceived and designed the experiments, performed the experiments, analysed the data, prepared figures and/or tables, authored or reviewed drafts of the paper, and approved the final draft.
- Ali Movasati analysed the data, prepared figures and/or tables, and approved the final draft.
- Véronique Clerc performed the experiments, analysed the data, prepared figures and/or tables, and approved the final draft.
- Jurek Andres performed the experiments, analysed the data, prepared figures and/or tables, and approved the final draft.
- Dr. Adriana Hotz performed the experiments, prepared figures and/or tables, and approved the final draft.
- Dr. Daniela Ferreira Garcia Rodrigues performed the experiments and approved the final draft.
- Prof. Dr. Karin J. Metzner conceived and designed the experiments, analysed the data, authored or reviewed drafts of the paper, and approved the final draft.

## Supplementary information

Supplementary Table S1. List of gRNAs present in the CRISPRko DDR library.

Supplementary Table S2. Summary of the sequencing results of the CRISPR/Cas9 DNA damage response (DDR) screening.

Supplementary Table S3. MAGeCK scores for CRISPR/Cas9 DNA damage response (DDR) screening replicates.

Supplementary Table S4. Statistical analysis for Figure 4.

Supplementary Table S5. Sequences of gRNAs used for CRISPR activation.

Supplementary Table S6. PCR/qPCR primers and cycling conditions.

Supplementary Figure S1. CRISPR/Cas9 DNA damage response (DDR) screening identifies genes potentially involved in the formation of large internal deletions (LID) in proviruses.

Supplementary Figure S2. Validation of the CRISPR/Cas9 DNA damage response (DDR) screening.

Supplementary Figure S3. Expression of the candidate genes involved in the formation of large internal deletions (LID) in proviruses.

Supplementary Figure S4. Validation of the CRISPRa-mediated overexpression of the candidate genes involved in the formation of large internal deletions (LID) in proviruses.

Supplementary Figure S5. Validation of the siRNA-mediated silencing of the candidate genes involved in the formation of large internal deletions (LID) in proviruses.

## Additional information and declaration

The manuscript’s text was edited for grammar and clarity using an AI-based language tool (Perplexity); all scientific content and interpretations are the authors’ own.

## References

1. Barré-Sinoussi, F., et al., Isolation of a T-lymphotropic retrovirus from a patient at risk for acquired immune deficiency syndrome (AIDS). Science, 1983. 220(4599).

2. World Health Organization. HIV and AIDS 2025 July 15, 2025]; Available from: https://www.who.int/news-room/fact-sheets/detail/hiv-aids.

3. Moir, S., T.W. Chun, and A.S. Fauci, Pathogenic mechanisms of HIV disease. Annu Rev Pathol, 2011. 6: p. 223–48.

4. Deeks, S.G., et al., HIV infection. Nat Rev Dis Primers, 2015. 1: p. 15035.

5. Craigie, R. and F.D. Bushman, HIV DNA integration. Cold Spring Harb Perspect Med, 2012. 2(7): p. a006890.

6. Skalka, A.M. and R.A. Katz, Retroviral DNA integration and the DNA damage response. Cell Death Differ, 2005. 12 **Suppl 1**: p. 971–8.

7. Knyazhanskaya, E., et al., NHEJ pathway is involved in post-integrational DNA repair due to Ku70 binding to HIV-1 integrase. Retrovirology, 2019. 16(1): p. 30.

8. Espeseth, A.S., et al., siRNA screening of a targeted library of DNA repair factors in HIV infection reveals a role for base excision repair in HIV integration. PLoS One, 2011. 6(3): p. e17612.

9. Fu, S., et al., HIV-1 exploits the Fanconi anemia pathway for viral DNA integration. Cell Rep, 2022. 39(8): p. 110840.

10. Bruner, K.M., et al., Defective proviruses rapidly accumulate during acute HIV-1 infection. Nat Med, 2016. 22(9): p. 1043–9.

11. Pollack, R.A., et al., Defective HIV-1 Proviruses Are Expressed and Can Be Recognized by Cytotoxic T Lymphocytes, which Shape the Proviral Landscape. Cell Host Microbe, 2017. 21(4): p. 494–506 e4.

12. Sanchez, G., et al., Accumulation of defective viral genomes in peripheral blood mononuclear cells of human immunodeficiency virus type 1-infected individuals. J Virol, 1997. 71(3): p. 2233–2240.

13. Ho, Y.C., et al., Replication-competent noninduced proviruses in the latent reservoir increase barrier to HIV-1 cure. Cell, 2013. 155(3): p. 540–51.

14. Pathakab, V.K. and W.-S. Hu, “Might as Well Jump!” Template Switching by Retroviral Reverse Transcriptase, Defective Genome Formation, and Recombination. Seminars in Virology, 1997. 8(2): p. 141–150.

15. Pulsinelli, G.A. and H.M. Temin, Characterization of large deletions occurring during a single round of retrovirus vector replication: novel deletion mechanism involving errors in strand transfer. J Virol, 1991. 65(9).

16. Imamichi, H., et al., Defective HIV-1 proviruses produce novel protein-coding RNA species in HIV-infected patients on combination antiretroviral therapy. Proc Natl Acad Sci U S A, 2016. 113(31): p. 8783–8.

17. Inderbitzin, A., et al., HIV-1 promoter is gradually silenced when integrated into BACH2 in Jurkat T-cells. PeerJ, 2020. 8: p. e10321.

18. Kok, Y.L., et al., Spontaneous reactivation of latent HIV-1 promoters is linked to the cell cycle as revealed by a genetic-insulators-containing dual-fluorescence HIV-1-based vector. Sci Rep, 2018. 8(1): p. 10204.

19. Su, D., et al., CRISPR/CAS9-based DNA damage response screens reveal gene-drug interactions. DNA Repair (Amst), 2020. 87: p. 102803.

20. Li, W., et al., MAGeCK enables robust identification of essential genes from genome-scale CRISPR/Cas9 knockout screens. Genome Biol, 2014. 15(554).

21. Inderbitzin, A., et al., Quantification of transgene expression in GSH AAVS1 with a novel CRISPR/Cas9-based approach reveals high transcriptional variation. Molecular Therapy - Methods & Clinical Development, 2022. 26: p. 107–118.

22. Li, W., et al., Quality control, modeling, and visualization of CRISPR screens with MAGeCK-VISPR. Genome Biol, 2015. 16(281).

23. Li, l., et al., Role of the non-homologous DNA end joining pathway in the early steps of retroviral infection. EMBO J., 2001. 20(12): p. 3272–81.

24. Zheng, Y., et al., Host protein Ku70 binds and protects HIV-1 integrase from proteasomal degradation and is required for HIV replication. J Biol Chem, 2011. 286(20): p. 17722–35.

25. Lahouassa, H., et al., HIV-1 Vpr degrades the HLTF DNA translocase in T cells and macrophages. Proc Natl Acad Sci U S A, 2016. 113(19): p. 5311–6.

26. Hrecka, K., et al., Lentiviral Vpr usurps Cul4-DDB1[VprBP] E3 ubiquitin ligase to modulate cell cycle. PNAS, 2007. 104(28): p. 11778–83.

27. Yan, J., et al., HIV-1 Vpr counteracts HLTF-mediated restriction of HIV-1 infection in T cells. Proc Natl Acad Sci U S A, 2019. 116(19): p. 9568–9577.

28. Cosnefroy, O., et al., Stimulation of the human RAD51 nucleofilament restricts HIV-1 integration in vitro and in infected cells. J Virol, 2012. 86(1): p. 513–26.

29. Shun, M.C., et al., LEDGF/p75 functions downstream from preintegration complex formation to effect gene-specific HIV-1 integration. Genes Dev, 2007. 21(14): p. 1767–78.

30. Mulder, L.C., L.A. Chakrabarti, and M.A. Muesing, Interaction of HIV-1 integrase with DNA repair protein hRad18. J Biol Chem, 2002. 277(30): p. 27489–93.

31. Golumbeanu, M., et al., Proteo-Transcriptomic Dynamics of Cellular Response to HIV-1 Infection. Sci Rep, 2019. 9(1): p. 213.

32. Chavez, A., et al., Highly efficient Cas9-mediated transcriptional programming. Nat Methods, 2015. 12(4): p. 326–8.

33. Sanson, K.R., et al., Optimized libraries for CRISPR-Cas9 genetic screens with multiple modalities. Nat Commun, 2018. 9(1): p. 5416.

34. Konermann, S., et al., Genome-scale transcriptional activation by an engineered CRISPR-Cas9 complex. Nature, 2015. 517(7536): p. 583–8.

35. Cui, Z., et al., MYO1F regulates T-cell activation and glycolytic metabolism by promoting the acetylation of GAPDH. Cell Mol Immunol, 2025. 22(2): p. 176–190.

36. Lada, S.M., et al., Quantitation of Integrated HIV Provirus by Pulsed-Field Gel Electrophoresis and Droplet Digital PCR. J Clin Microbiol, 2018. 56(12).

37. Strouvelle, V.P., et al., No Effect of Pegylated Interferon-alpha on Total HIV-1 DNA Load in HIV-1/HCV Coinfected Patients. J Infect Dis, 2018. 217(12): p. 1883–1888.

38. Schrijvers, R., et al., LEDGF/p75-independent HIV-1 replication demonstrates a role for HRP-2 and remains sensitive to inhibition by LEDGINs. PLoS Pathog, 2012. 8(3): p. e1002558.

39. Kile, A.C., et al., HLTF’s Ancient HIRAN Domain Binds 3’ DNA Ends to Drive Replication Fork Reversal. Mol Cell, 2015. 58(6): p. 1090–100.

40. Bai, G., et al., HLTF Promotes Fork Reversal, Limiting Replication Stress Resistance and Preventing Multiple Mechanisms of Unrestrained DNA Synthesis. Mol Cell, 2020. 78(6): p. 1237–1251 e7.

41. Sheridan, P.L., et al., Cloning of an SNF2/SWI2-related protein that binds specifically to the SPH motifs of the SV40 enhancer and to the HIV-1 promoter. J Biol Chem, 1995. 270(9): p. 4575–87.

42. Ding, H., et al., Characterization of a helicase-like transcription factor involved in the expression of the human plasminogen activator inhibitor-1 gene. DNA Cell Biol, 1996. 15(6): p. 429–42.

43. Nightingale, K., et al., High-Definition Analysis of Host Protein Stability during Human Cytomegalovirus Infection Reveals Antiviral Factors and Viral Evasion Mechanisms. Cell Host Microbe, 2018. 24(3): p. 447–460 e11.

44. Panda, S.K., et al., IL-4 controls activated neutrophil FcgammaR2b expression and migration into inflamed joints. Proc Natl Acad Sci U S A, 2020. 117(6): p. 3103–3113.

45. Ablashi, D.V., et al., Human herpesvirus-7 (HHV-7): current status. Clinical and Diagnostic Virology, 1995. 4: p. 1–13.

46. Fehse, B., et al., Pois(s)on--it’s a question of dose. Gene Ther, 2004. 11(11): p. 879–81.

47. Hatakeyama, M., et al., SUSHI: an exquisite recipe for fully documented, reproducible and reusable NGS data analysis. BMC Bioinformatics, 2016. 17(1): p. 228.

48. Shalem, O., et al., Genome-scale CRISPR-Cas9 knockout screening in human cells. Science, 2014. 343(6166): p. 84–87.

