## Supplementary figures for "DNA Damage Response Proteins Are Involved in the Formation of Defective HIV-1 Proviruses"

**Supplementary Table S1. List of gRNAs present in the CRISPRko DDR library.**

The CRISPRko DDR library has been published by Su et al., 2020.

**Supplementary Table S2. Summary of the sequencing results of the CRISPR/Cas9 DNA damage response (DDR) screening.**

Generated using MAGeCK Count Report. DDR 1-3, biological replicates of the CRISPR/Cas9 screening; NGS 1-3, next generation sequencing runs of the CRISPR/Cas9 screening.

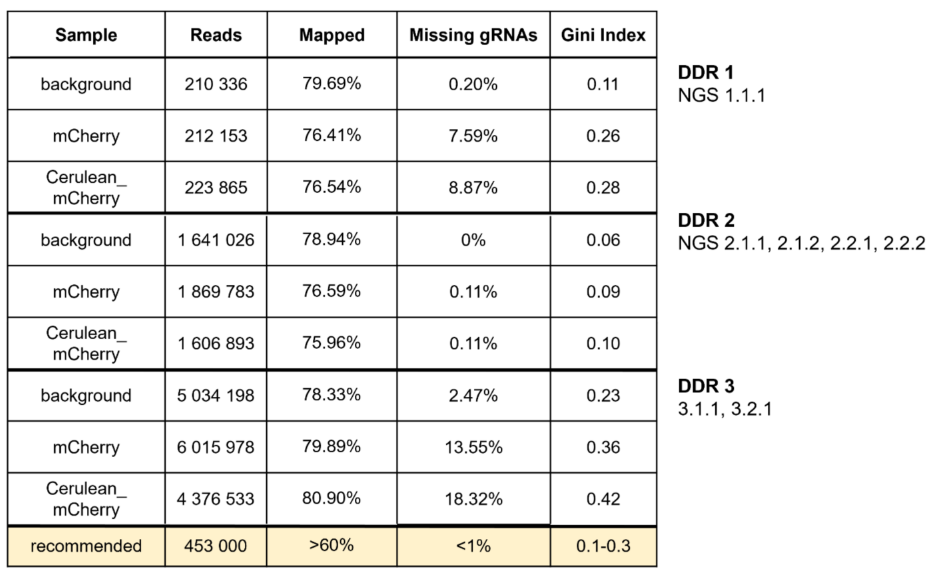

**Supplementary Table S3. MAGeCK scores for CRISPR/Cas9 DNA damage response (DDR) screening replicates.**

The gRNA counts in mCherry+ and Cerulean+/mCherry+ (Dp) samples were compared using MAGeCK for the NGS 1.1.1, NGS 2.1.1, 2.1.2, 2.2.1, 2.2.2, and NGS 3.1.1, 3.2.1. Additionally, combined comparisons were made for all NGS runs from the second replicate (NGS 2) and third screen replicate (NGS 3).

**Supplementary Table S4. Statistical analysis for Figure 4.**

**Supplementary Table S5. Sequences of gRNAs used for CRISPR activation.**

**Supplementary Table S6. PCR/qPCR primers and cycling conditions.**

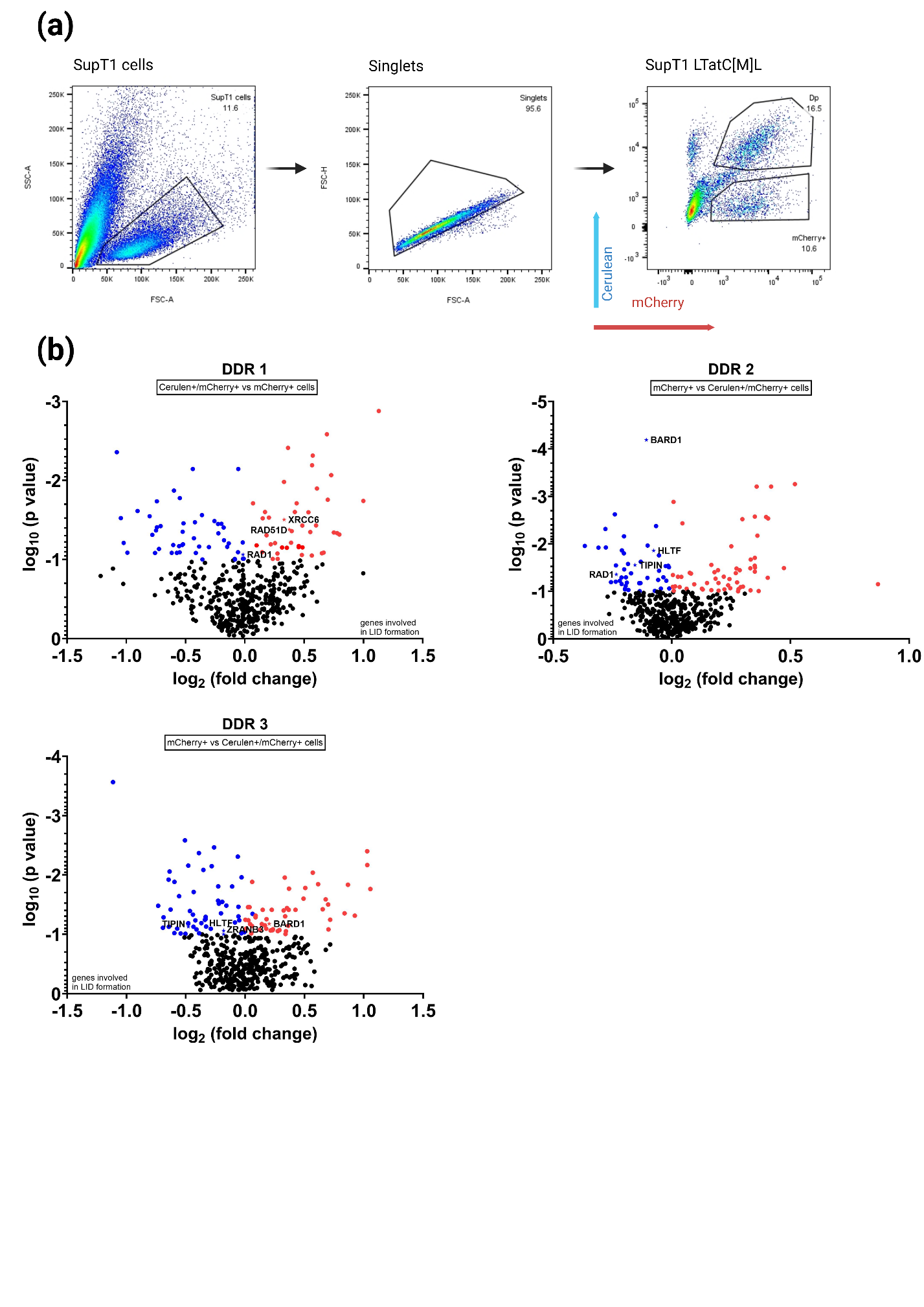

**Supplementary Figure S1.** **CRISPR/Cas9 DNA damage response (DDR) screening identifies genes potentially involved in the formation of large internal deletions (LID) in proviruses.**

(a) Flow cytometry gating strategy for sorting of the SupT1 cells transduced with CRISPRko DDR library and LTatC[M]L vector. Example shown for the second screen CRISPR/Cas9 DDR screening replicate (DDR 2). (b) Volcano plots showing MAGeCK gene enrichment scores and associated gRNA log2 fold changes for CRISPR/Cas9 screen replicates. Significantly enriched and depleted gRNA (adjusted p value < 0.1) are coloured in red and blue, respectively; selected candidate genes are marked with a star.

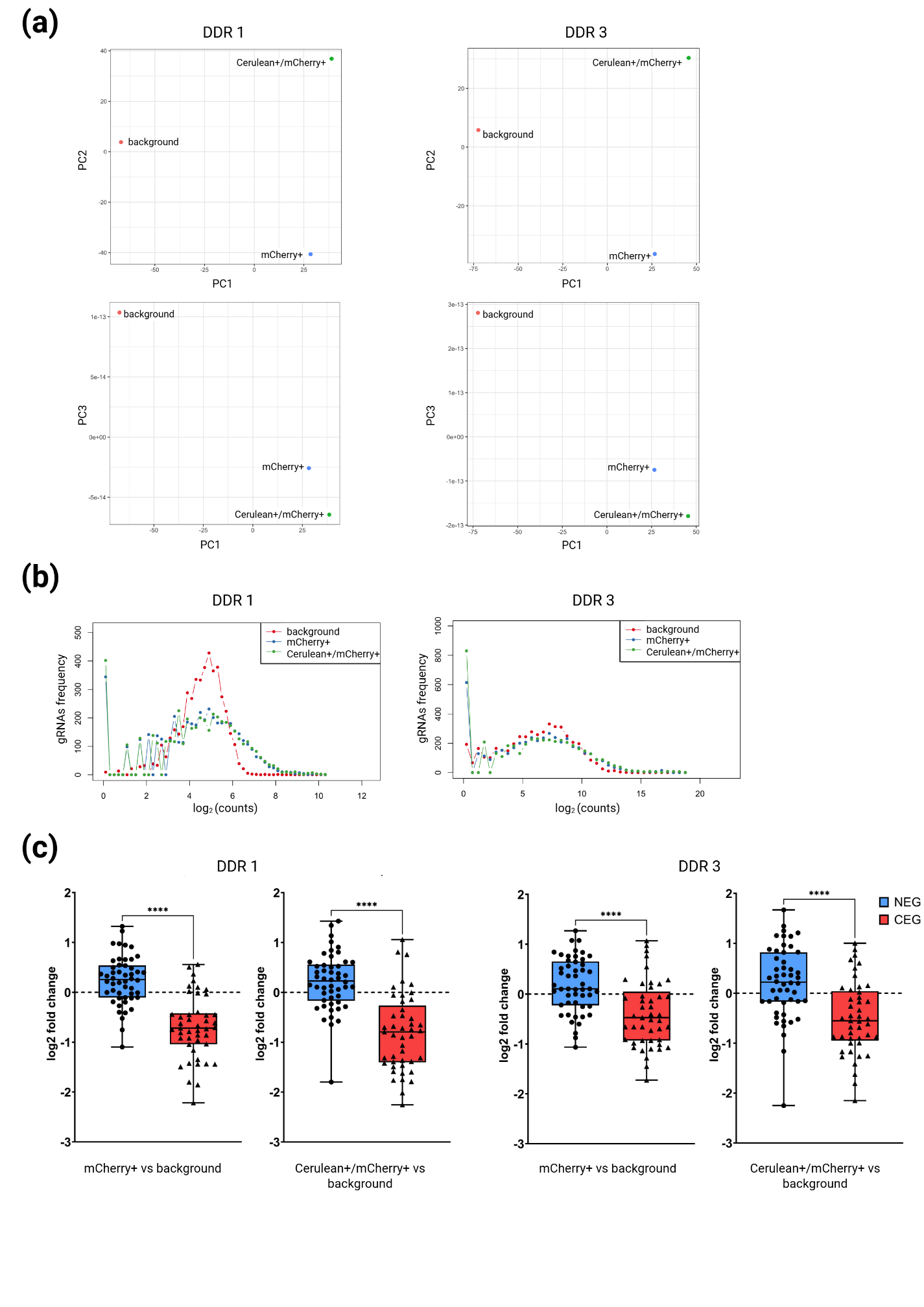

**Supplementary Figure S2. Validation of the CRISPR/Cas9 DNA damage response (DDR) screening.**

(a) Principal component analysis (PCA) of the double-positive Cerulean+/mCherry+ cells, single mCherry+ cells, and background control samples. Plots shown for the first (DDR 1) and the third (DDR 3) CRISPR/Cas9 screening replicates. (b) gRNAs read count distribution in double-positive Cerulean+/mCherry+ cells, the single mCherry+ cells, and background control samples. C) Internal controls of the CRISPR/Cas9 screening: log fold change in the count of gRNAs targeting core-essential (CEG) and non-essential genes (NEG) during CRISPR/Cas9 screening.  Data was statistically tested using a Mann-Whitney U-Test, p value <0.05 was considered significant, **** = p <0.0001.

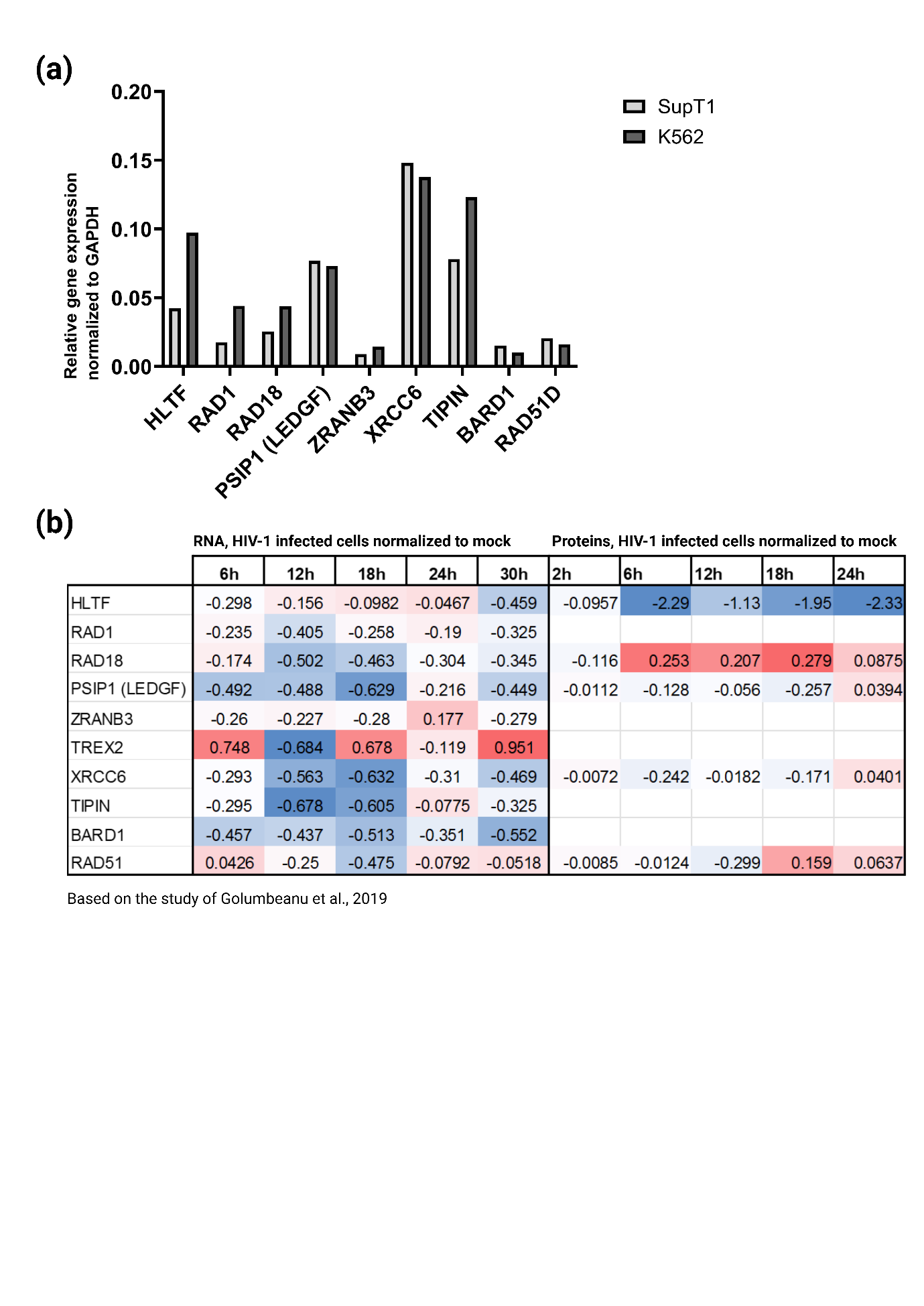

**Supplementary Figure S3. Expression of the candidate genes involved in the formation of large internal deletions (LID) in proviruses.**

(a) Baseline gene expression of candidate genes in the SupT1 and K562 cells, normalized to *GAPDH*. (b) Expression of candidate genes at different time points during HIV-1eGFP infection in the SupT1 CD4+ cells, from the left: measured on the RNA and protein level. The data is colour-coded based on the expression level, with the lowest expression marked in blue and the highest in red. The table was compiled based on the data available through PEACHi2 online resource (Golumbeanu et al., 2019).

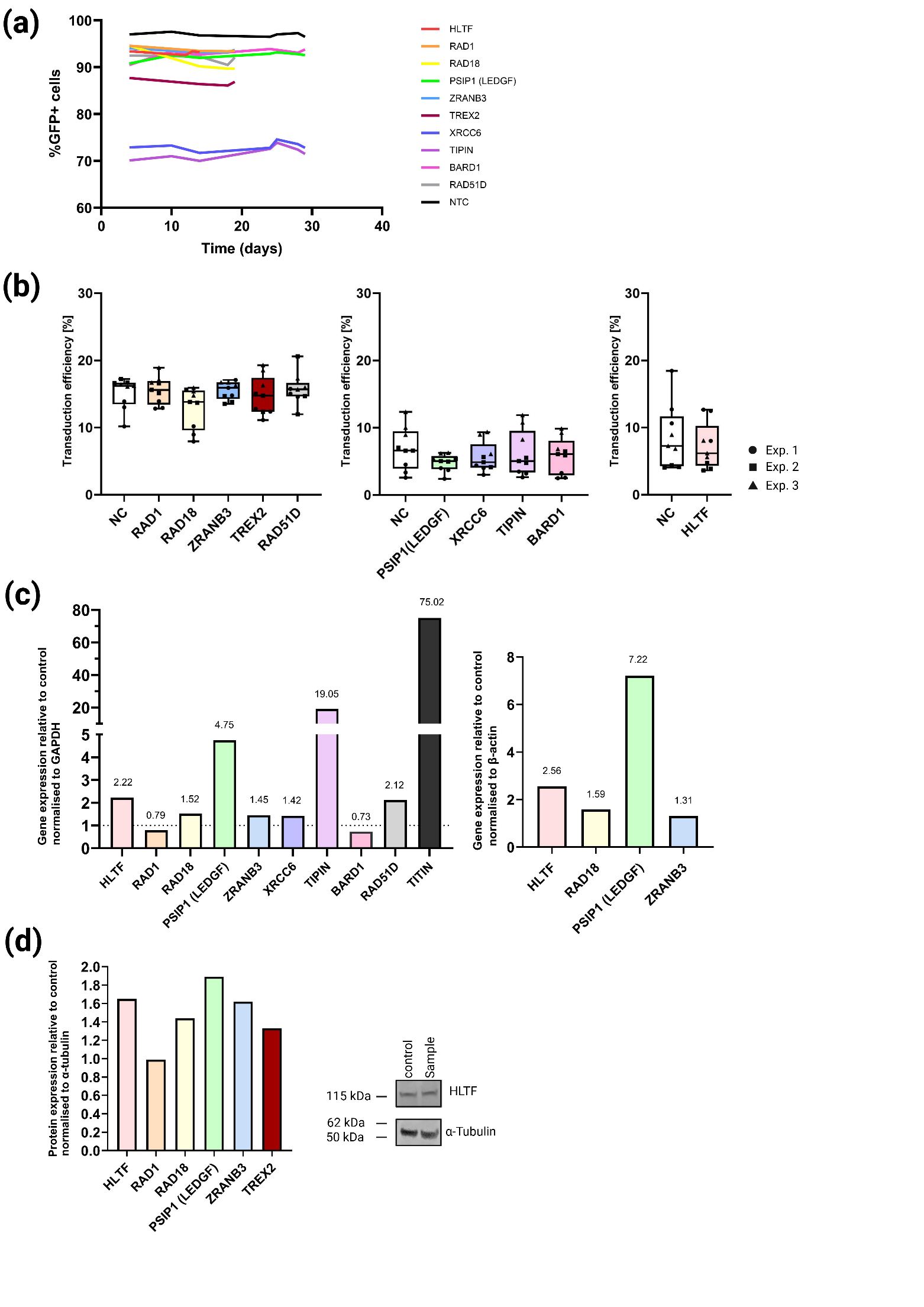

**Supplementary Figure S4. Validation of the CRISPRa-mediated overexpression of the candidate genes involved in the formation of large internal deletions (LID) in proviruses.**

(a) Percentage of K562-VPR cells transduced with CRISPRa-gRNA vector (GFP+) present in cell culture over time, as measured by flow cytometry. (b) Transduction efficiency of LTatC[M]L vector. Each data point represents one experiment, corresponding to Figure 4b. (c) Expression of candidate genes in K562-VPR cells transduced with corresponding CRISPRa-gRNA vector, measured by RT-qPCR. Data shown relative to non-targeting control gRNA, normalized to two housekeeping genes, from the left: *GAPDH* and *β-actin*. (d) Quantification of protein expression of candidate genes in K562 VPR cells transduced with corresponding CRISPRa-gRNA vector, measured by Western blot, example for HLTF protein expression shown on the right Data shown relative to non-targeting control gRNA, normalized to α-tubulin.

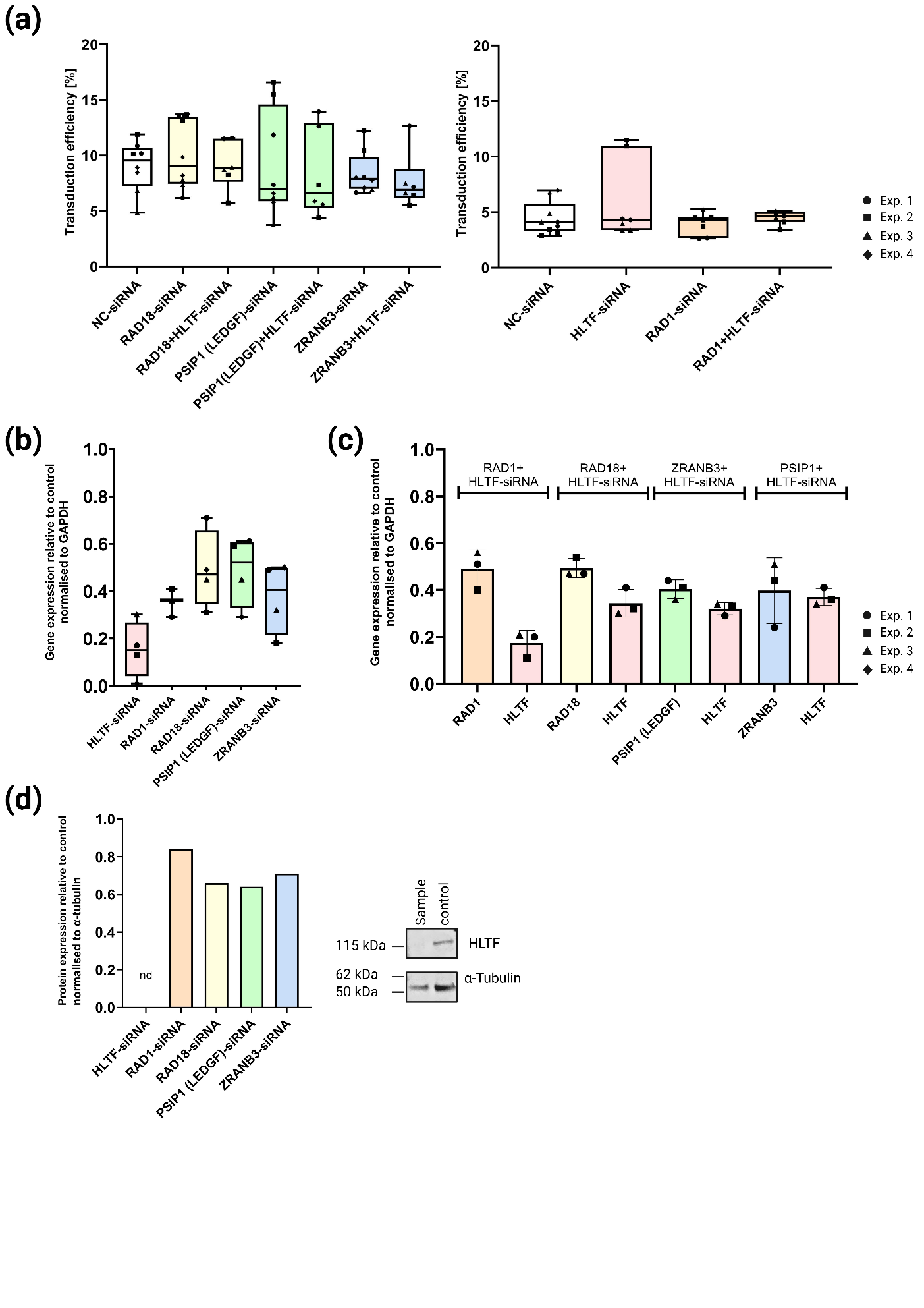

**Supplementary Figure S5. Validation of the siRNA-mediated silencing of the candidate genes involved in the formation of large internal deletions (LID) in proviruses.**

(a) Transduction efficiency of LTatC[M]L vector. Each data point represents one experiment, corresponding to Figure 4c. (b) Expression of candidate genes in K562 cells transfected with corresponding siRNA, measured by RT-qPCR. Data shown relative to scrambled control siRNA, normalized to *GAPDH*. Each data point represents one experiment, corresponding to Figure 4c. (c) Expression of candidate genes in K562 cells transfected with a combination of siRNAs targeting two genes, measured by RT-qPCR. Data shown relative to scrambled control siRNA, normalized to *GAPDH*. Each data point represents one experiment, corresponding to Figure 4c. (d) Quantification of protein expression of candidate genes in K562 cells transfected with corresponding siRNA, measured by Western blot, example for HLTF protein expression shown on the right (nd, not detected). Data shown relative to control scrambled siRNA, normalized to α-tubulin.
