## Supplementary table 6 for "DNA Damage Response Proteins Are Involved in the Formation of Defective HIV-1 Proviruses"

| **Target** | **Name** | **Sequence (5’-3’)** | **Annealing temperature (°C)** | **Elongation time (sec)** | **Amplicon length (bp)** |
| --- | --- | --- | --- | --- | --- |
| RAD1 | RAD1_748 | GAAGTTCCCACCTTGACTATCC | 56 | 40 | 546 |
|  | RAD1_1393rc | CTTGAAATGTTACGCTGACTCAC |  |  |  |
| HLTF | HLTF_879 | CATGAAATGGAACCAGCTGAGG | 57 | 40 | 479 |
|  | HLTF_1357rc | CTTTTGGGGCGGGAGCTAGA |  |  |  |
| RAD18 | RAD18_87 | TAGAGTCACCAGCCAAATCTC | 53 | 35 | 482 |
|  | RAD18_568rc | CTTTAAATCACGATCAGAGAGC |  |  |  |
| TREX2 | TREX2_265 | TAGTATTGCCCCGGGTCCTG | 57 | 35 | 543 |
|  | TREX2_807rc | GGTCATCAGGCGGCAAGTAC |  |  |  |
| BARD1 | BARD1_285 | TCTGCGGCCTGTCGATTATAC | 54 | 25 | 303 |
|  | BARD1_587rc | CATCCATTGAGAATCCCAAGC |  |  |  |
| TIPIN | TIPIN_337 | AGACTTGAAGATGCTAATCAGAC | 55 | 25 | 360 |
|  | TIPIN_696rc | AGCTTTGCCTGCCTTCTTTCC |  |  |  |
| TIPIN | TIPIN_2 | TGCTAATCAGACACATGGAGC | 54 | 25 | 504 |
|  | TIPIN_2rc | TGGCAATAGCATCATTACATGG |  |  |  |
| RAD51D | RAD51D_553 | CAACCACATAACTCGAGACAG | 55 | 25 | 399 |
|  | RAD51D_951rc | GAGAGTGAGGCCAAGGAAC |  |  |  |
| ZRANB3 | ZRANB3_1700 | AGTTGATGACCTCCTGTGAC | 53 | 35 | 483 |
|  | ZRANB3_2182rc | CAGGTTCTGACTGTGCAAGG |  |  |  |
| XRCC6 (Ku70) | XRCC6_480 | CTAGAGCTTGACCAGTTTAAGG | 55 | 25 | 414 |
|  | XRCC6_893rc | TCGCTTCCTGGTCTCCTTG |  |  |  |
| XRCC6 (Ku70) | XRCC6_2 | ACATGATGGGCCACGGATC | 55 | 25 | 537 |
|  | XRCC6_2rc | GAAGCAAACCGCCTGTACTTG |  |  |  |
| LEDGF | LEDGF_974 | CCTAAAAAGCAGCCTAAGAAGG | 52 | 20 | 295 |
|  | LEDGF_1268rc | CAGTTTCCATTTGTTCCTCTTG |  |  |  |
| TTN | TTN_6606 | GTACTGGCCCGAAGACAATG | 55 | 25 | 387 |
|  | TTN_6983rc | TCATCTTCCACAAGTACACAGC |  |  |  |
| β-actin | bact532 | CCATCTACGAGGGGTATGC | 55 | 30 | 196 |
|  | bact728rc | CGTGGCCATCTCTTGCTC |  |  |  |

Table 6.1. PCR primers and conditions for generating standards for qPCR

| 94°C | 3 min |  |
| --- | --- | --- |
| 94°C | 15 sec | 35x |
| 52-57°C | 30 sec |  |
| 72°C | 25-45 sec |  |
| 72°C | 10 min |  |
| 10°C | hold |  |

Each PCR reaction contained 2 μl of 10 x PCR Buffer (Sigma-Aldrich), 1.2 μl of 25 μM MgCl2 (Sigma-Aldrich), 0.4 μl of 10 μM dNTPs (Thermo Fisher Scientific), 0.4 μl of 20 μM forward primer, 0.4 μl of 20 μM reverse primer, and 0.2 μl JumpStart *Taq* DNA Polymerase (Sigma-Aldrich). The reaction was adjusted to a final volume of 18 μl with nuclease-free water. For each reaction, 2 μl of cDNA were added. Samples were run alongside the NO RT and PBS controls.

Table 6.2. qPCR primers and conditions

| **Target** | **Name** | **Sequence (5’-3’)** | **Annealing temperature (°C)** |
| --- | --- | --- | --- |
| GAPDH | mf45 | TCGACAGTCAGCCGCATCTT | 55 |
|  | mf46 | GGCAACAATATCCACTTTACCAG |  |
| RAD1 | RAD1_789 | GAAGCATTTCATTGTAATCAGACC | 55 |
|  | RAD1_910rc | AGGAAGCCTCTGTTATCTGTCC |  |
| HLTF | HLTF_938 | TCTAGCTTGGATGGTGTCACG | 55 |
|  | HLTF_1058rc | GACATTTTCTGGTCGGTCCTT |  |
| RAD18 | RAD18_qFw | AGCAGGGGAGCAGGTTAATG | 56 |
|  | RAD18_qRev | AGCCTCTGAGGGATCTGGAG |  |
| TREX2 | TREX2_379 | GGAAGGCTGGCTTTGATGGC | 57 |
|  | TREX2_499rc | GCACACAGCAGGGGGAAATC |  |
| BARD1 | BARD1_qFw | ATCCTCAGCTAGCCACTGC | 55 |
|  | BARD1_qRev | TTCTGAAGACAGCCCACTGC |  |
| TIPIN | TIPIN_qFw | ATGGAGCACTGGGCACATAG | 55 |
|  | TIPIN_qRev | TCCCAGGTATTCAACTCTGTC |  |
| RAD51D | RAD51D_qFw | TGTCTGGCCAAATCTTCCCG | 55 |
|  | RAD51D_qRev | AGGTGGCAGTAAACAGCAGG |  |
| ZRANB3 | ZRANB3_qFw | ATTGGCTGCCTCGGAAGACC | 58 |
|  | ZRANB3_qRev | CAGCCCTCTACAGGAAAGGC |  |
| XRCC6 (Ku70) | XRCC6_qFw | GGGTCTGTGCCAACCTCT | 53 |
|  | XRCC6_qRev | AAAGCCCCCAGGTTTCTTC |  |
| LEDGF | LEDGF_qFw | AGGGGTTACTTCAACCTCCG | 54 |
|  | LEDGF_qRev | TTGCGTTTTCGATCTGCTGC |  |
| TTN | TTN_qFw | GAGAAACCTCCAGTCACGC | 55 |
|  | TTN_qRev | CCCTCATGAACCTCCATACC |  |
| β-actin | bact532 | CCATCTACGAGGGGTATGC | 54 |
|  | bact728rc | CGTGGCCATCTCTTGCTC |  |

When multiple genes were analysed on the same qPCR plate, all reaction were run using the lowest annealing temperature of the corresponding primer sets.

| 94°C | 3 min |  |
| --- | --- | --- |
| 94°C | 15 sec | 50x |
| 53-58°C | 30 sec |  |
| 72°C | 30 sec |  |
| 95°C | 15 sec |  |
| 55°C | 1 min | melting curve |
| 95°C | 20 sec |  |

Table 6.3. qPCR of HIV-1 *pol* gene to assess the efficiency of BluePippin size selection

| **Target** | **Name** | **Sequence (5’-3’)** | **Annealing temperature (°C)** |
| --- | --- | --- | --- |
| CCR5 | mf51 | TCATTACACCTGCAGCTCTCATTTTCCATACAGTC | 60 |
|  | mf52 | CACCGAAGCAGAGTTTTTAGGATTCCCGAGTA |  |
|  | mf73tq probe | FAM-TGCCGCTGCTTGTCATGGTCATCTG-TAMRA |  |
| Pol | mf299 | GCACTTTAAATTTTCCCATTAGTCCTA | 55 |
|  | mf302 | CAAATTTCTACTAATGCTTTTATTTTTTC |  |
|  | mf348tq probe | FAM-AAGCCAGGAATGGATGGCC-TAMRA |  |
| 2 LTR | nef-U39146rc-w | GTGTGGTAGAYCCACAGATCAAG | 55 |
|  | 2LTR-fw | GCTTGCCTTGAGTGCTTCAAG |  |
|  | ca20tq | FAM-TGTGTGCCCGT-BHQ-1 |  |

| w for wobble |
| --- |
| underlined bases are locked nucleic acids |

Each 20 μl reaction comprised 10 μl of sample or standard and 10 μl of the following master mix: nuclease‑free H2O, 1 x Sigma PCR buffer (10 x stock, without MgCl₂), 3 mM MgCl2, 0.4 mM of dNTPs, 2.5 nM ROX passive reference dye, 0.3 μM dual‐labelled probe, 1 μM forward primer, 1 μM reverse primer and 0.5 U JumpStart Taq DNA Polymerase. Thermal cycling was performed as described below.

| 95°C | 5 min |  |
| --- | --- | --- |
| 95°C | 15 sec | 50x |
| 55-60°C | 30 sec |  |
| 60°C | 30 sec |  |

Table 6.4. PCR amplification to map the deletions

| **Target** | **Name** | **Sequence (5’-3’)** |
| --- | --- | --- |
| LTatC[M]L | KYL-mCherry_rc | GAAGGACAGCTTCAAGTAGTCG |
|  | KYL-5LTRwg Fw | GACAAGAGATCCTTGATCTGTGGATC |

| **Cycling conditions:** |  |  |
| --- | --- | --- |
| 94°C | 2min |  |
| 94°C | 30s | 35 cycles |
| 58°C | 30s |  |
| 65°C | 5min |  |
| 65°C | 10min |  |
| 10°C | hold |  |
